# Testing hypotheses about the microbiome using the linear decomposition model (LDM)

**DOI:** 10.1101/229831

**Authors:** Yi-Juan Hu, Glen A. Satten

## Abstract

**Motivation:** Methods for analyzing microbiome data generally fall into one of two groups: tests of the global hypothesis of any microbiome effect, which do not provide any information on the contribution of individual operational taxonomic units (OTUs); and tests for individual OTUs, which do not typically provide a global test of microbiome effect. Without a unified approach, the findings of a global test may be hard to resolve with the findings at the individual OTU level. Further, many tests of individual OTU effects do not preserve the false discovery rate (FDR).

**Results:** We introduce the linear decomposition model (LDM), that provides a single analysis path that includes global tests of any effect of the microbiome, tests of the effects of individual OTUs while accounting for multiple testing by controlling the FDR, and a connection to distance-based ordination. The LDM accommodates both continuous and discrete variables (e.g., clinical outcomes, environmental factors) as well as interaction terms to be tested either singly or in combination, allows for adjustment of confounding covariates, and uses permutation-based *p*-values that can control for correlation. The LDM can also be applied to transformed data, and an “omnibus” test can easily combine results from analyses conducted on different transformation scales. We also provide a new implementation of PERMANOVA based on our approach. For global testing, our simulations indicate the LDM provided correct type I error and can have comparable power to existing distance-based methods. For testing individual OTUs, our simulations indicate the LDM controlled the FDR well. In contrast, DESeq2 often had inflated FDR; MetagenomeSeq generally had the lowest sensitivity. The flexibility of the LDM for a variety of microbiome studies is illustrated by the analysis of data from two microbiome studies. We also show that our implementation of PERMANOVA can outperform existing implementations.

## Background

Data from studies of the microbiome is accumulating at a rapid rate. The relative ease of conducting a census of bacteria by sequencing the 16S rRNA gene (or, for fungi, the 18S rRNA gene) has led to many studies that examine the association between microbiome and health states or outcomes. Many microbiome studies have complex design features (e.g., paired, clustered, or longitudinal data) or complexities that frequently arise in medical studies (e.g., the presence of confounding covariates). While a number of new methods for analyzing microbiome data have been recently proposed, there is still a need for methods that can account for the special features of microbiome data and the complex study designs in common use while preserving size or controlling the false discovery rate (FDR).

Statistical methods for analyzing microbiome data seem to fall into one of two camps. One camp comprises methods that test the *global* effect of the microbiome, such as PERMANOVA [Anderson, 2001, McArdle and Anderson, 2001], MiRKAT [Zhao *et al.*, 2015], aMiSPU [Wu *et al.*, 2016], and pairNM [Shi and Li, 2017], which can be used to test the hypothesis that variables of interest (e.g., case-control status) are significantly associated with overall microbial compositions. However, these methods do not provide convenient tests of the effects or contributions of individual operational taxonomic units (OTUs), should a global microbiome effect be found (here we use “OTU” generically to refer to any feature such as amplicon sequence variants or any other taxonomic or functional grouping of bacterial sequences). The other camp is comprised of OTU-by-OTU tests, often directly using a method developed for RNA-Seq data such as DESeq2 [Love *et al.*, 2014] and edgeR [Robinson *et al.*, 2010] or a modification thereof such as metagenomeSeq [Paulson *et al.*, 2013]; some other methods in this camp have adopted a compositional data approach (such as ANCOM [Kaul *et al.*, 2017, Mandal *et al.*, 2015] and ALDEx2 [Fernandes *et al.*, 2014]), were developed for longitudinal data (such as ZIBR [Chen and Li, 2016]), or employed a multi-stage strategy (such as massMap [Hu *et al.*, 2018]). While some of these approaches have been widely applied, they generally do not give a single test of the global null hypothesis. Although test statistics or *p*-values from OTU-specific tests can of course be combined to give a global test, the performance of this kind of global test is often poor since many of the OTU-specific tests only contribute noise.

We introduce here the Linear Decomposition Model (LDM) for analyzing microbial count or relative abundance data that are obtained in a 16S rRNA study or a shotgun metagenomics sequencing study. The LDM gives a unified approach that allows both global testing of the overall effect of the microbiome on arbitrary traits of interest, while also providing OTU-specific tests that correspond to the contribution of individual OTUs to the global test results. It allows for complex fixed-effects models such as models that include multiple variables of interest (both continuous and categorical), their interactions, as well as confounding covariates. It is permutation based, and so can accommodate clustered data and maintain validity for small sample sizes and when data are subject to overdispersion. Because the permutations are based on the Freedman-Lane approach [Freedman and Lane, 1983], we can construct powerful type III or “last variable added” tests like those used in most linear regression packages [Kleinbaum *et al.*, 2007, Muller and Fetterman, 2012]. We also provide a new version of the PERMANOVA test based on our approach that we show outperforms the functions `adonis` and `adonis2` in the *R* package `vegan`, the most commonly used implementations of PERMANOVA. Recent simulation studies suggest that many microbiome analysis methods fail to control the FDR when applied to overdispersed data [Hawinkel *et al.*, 2017]. We show that the LDM controls FDR in exactly the kind of situations where other methods fail.

We describe the LDM in detail in the methods section. In the results section, we describe the simulation studies and the two real datasets that we use to assess the performance of the LDM, and compare it to results obtained by PERMANOVA, MiRKAT, aMiSPU, DESeq2, edgeR, MetagenomeSeq, the Wilcoxon rank-sum test, ANCOM, and ALDEx2. We conclude with a discussion section. Some technical details are relegated to Supplementary Materials.

## Methods

Data from a microbiome study are usually summarized in a table of read counts, here referred to as the OTU table and denoted by *Y*. The OTU table is the *n* × *J* data matrix whose (*i, j*)^th^ element is the number of times OTU *j* is observed in sample *i*. The total counts in each sample (the library size) can vary widely between samples and this variability must be accounted for. Here we accomplish this by converting counts to frequencies (i.e., relative abundances) by dividing the raw counts in each sample by the library size of that sample, although other normalizations can be used with the LDM if desired. Our simulation studies showed that the LDM performs well when counts are scaled to frequencies, even with highly overdispersed data. Although the LDM accounts for the compositionality of the frequency data by using an appropriate permutation scheme, a compositional analysis in the sense of Aitchison [1986] can also be conducted by applying the LDM to appropriately transformed (e.g., centered log-ratio) data.

### The LDM as a Linear Model

Many reasonable models of the relationship between data in an OTU table and covariates that describe traits or characteristics of individual samples can be expressed as a linear model. Because the large number of OTUs is a problem for models in which OTU frequencies predict traits or covariates when the goal is inference, we consider models in which traits and covariates are used as predictors of OTU frequencies. Thus, we consider the model

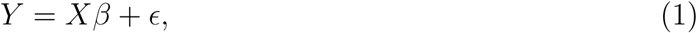

where *X* is an *n* × *r* design matrix containing *r* covariates (including traits and confounders), *β* is an *r* × *J* matrix that we must estimate, and *ϵ* is an *n* × *J* matrix of error terms with E(*ϵ*|*X*) = 0. Considered as *J* models for the columns of *Y*, the *j*th column of *β* are the regression coefficients for the *j*th regression model. In order to provide a clean decomposition of the sum of squares of *Y*, we will require that the columns of *X* are *orthonormal*. This also aids in the interpretation of some hypothesis tests, particularly terms that represent interactions with main effect terms that are also being tested. A motivating example for (1) is when *X* contains only an intercept and an indicator for a binary trait such as case-control status; then the first row of *β* is proportional to the means of the OTU frequencies while the second row of *β* is proportional to the differences in the OTU frequencies between case and control participants.

The least-squares estimators for the matrix *β* can be obtained by minimizing 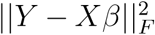 with respect to *β* (holding *X* and *Y* fixed), where 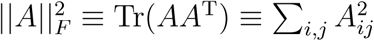 is the Frobenius (matrix) norm of matrix *A* and Tr(.) is the trace operator. Satten *et al.* [2017] showed that the resulting estimators are 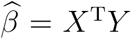, which are also the estimators obtained by fitting the regression model column by column. If we write 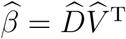 where 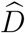 is a diagonal matrix with elements given by the norms of the rows of 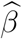, so that the columns of 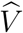 are normalized rows of 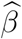, then we may rewrite (1) as

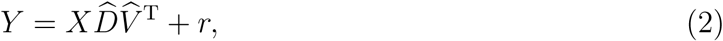

where 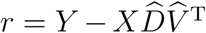 is the residual matrix. Equation (2) is the basic form of the LDM. It is similar to the singular value decomposition (SVD) in that a matrix *Y* is decomposed into the product of an orthogonal matrix *X*, a diagonal matrix *D* with positive elements and a third matrix *V*, but differs from the SVD because *X* may be chosen in any convenient way as long as it has orthonormal columns, and because the columns of *V* are not orthogonal.

In some situations, we may wish to partition covariates into *K* groups, which we call “submodels”. For example, we may wish to group several measures of smoking history into a single smoking “submodel”. In this case, we partition *X* as (*X*_1_, …, *X*_*K*_), and re-write (1) and (2) as

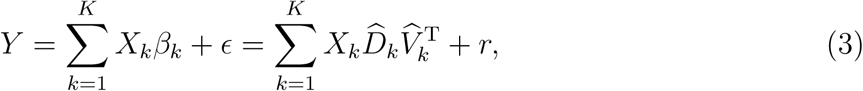

where the least-squares estimators of *β*_*k*_ are given by 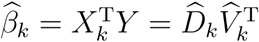 for each *k*. We will refer to the number of linearly-independent columns in *X*_*k*_ as the number of *components* in the submodel.

The LDM (3) can be used to describe the amount of the total sum of squares 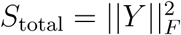 that is explained by each submodel *X*_*k*_ or the full model *X* by noting that 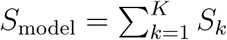 where 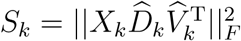 for *k* = 1, …, *K*. In analogy with Satten *et al.* [2017] we can express *S*_*k*_ in one of two ways:

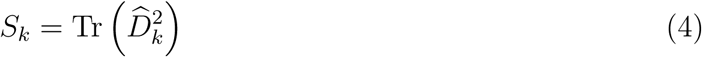

or

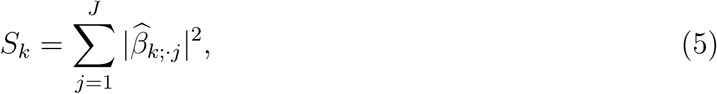

where 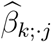 is the *j*th column of 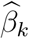 and |*b*| is the Euclidean norm of vector *b*. The first representation indicates that, as with a standard SVD, the sum of squares for the *k*th submodel is given by the sum of squares of the corresponding singular values (diagonal elements of 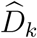). The second representation partitions *S*_*k*_ into contributions from each OTU. Note these results only require *X* has orthonormal columns, not 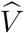.

### Testing hypotheses using the LDM

The LDM uses the decomposition of the sum of squares in (4) and (5) to test global and OTU-specific null hypotheses about the effect of individual covariates or sets of covariates grouped into submodels. To test hypotheses about the contribution of the *j*th OTU to the sum of squares for the *k*th submodel, we use its contribution to the model sum of squares given in (5), normalized as an *F* statistic, to give the test statistic

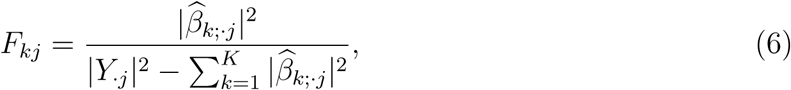

where *Y*_·*j*_ is the *j*th column of *Y* and we have dropped the constant of proportionality {Rank(*Y*) − Rank(*X*)} */*Rank(*X*_*k*_) found in a typical *F* test as we intend to use permutation to assess significance.

To test the global hypothesis we consider the total sum of squares for the *k*th submodel given in (4), again normalized as an *F* statistic. This statistic can either be constructed directly, or by summing the numerator and denominator of the OTU-specific statistics (5) separately, i.e.,

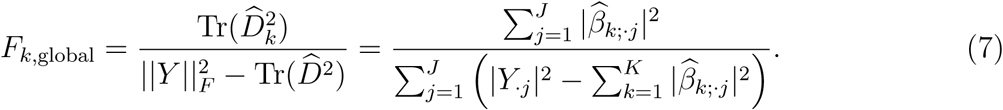

The presence of all submodels in the denominator indicates that this is a type III or “last variable added” test statistic. As before, the usual constant of proportionality in a *F*-statistic is dropped because significance will be assessed using permutation. Note that OTU-specific and global tests are linked through (6) and (7) in a natural way. If *Y* is centered, the sums-of-squares in (4) and (5) are proportional to the *variance* explained by a submodel or an individual OTU in a submodel; for this reason we sometimes refer to the *F*-tests in the LDM as tests of the variance explained (VE).

### The LDM as a decomposition

The LDM in equation (3) can also lead to an exact decomposition of the OTU table *Y*, if we augment *X* with extra columns *X*_*r*_ so that the full set of columns 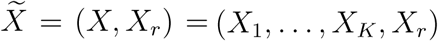 span the column space of *Y*. The “extra” columns *X*_*r*_ are used to decompose the residual matrix *r*. If we further write 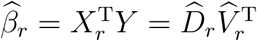, where 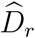 is a diagonal matrix with entries given by the norms of the rows of 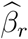, so that the columns of 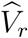 are normalized, then we can rewrite (3) as

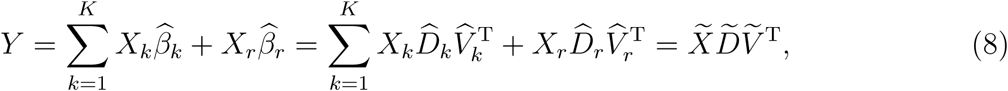

where 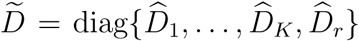 and 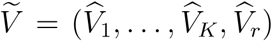. Thus the LDM, like the SVD, can give a full-rank decomposition of *Y*.

We can establish a connection between the LDM and distance-based methods such as PERMANOVA by choosing *X*_*r*_ to be the eigenvectors of a (residual) distance matrix Δ_*r*_ = (*I* − *XX*^T^)Δ(*I* − *XX*^T^) that have non-zero eigenvalues, where *XX*^T^ is the hat matrix of *X*. Note these eigenvectors are always orthogonal to the columns of *X*. This choice allows direct comparison of the variability explained by covariates to the variability explained by the principal components of a (residual) distance matrix, in analogy with PERMANOVA. A *scree plot* based on values of 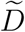 in (8) can visually indicate whether that values of 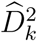 are “large” relative to the residual error. To help ensure that the rank of 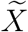 and Δ agree, if *Y* has been column-centered, then we also center Δ as recommended by Gower [1966]. Then, the columns of both 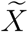 and *Y* are orthogonal to the vector 1. Some caution is necessary for submodels having more than one component; although *S*_*k*_ in (4) is invariant to the choice of basis for *X*_*k*_, the individual diagonal elements of 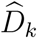 depend on the choice of basis. Thus, we recommend these elements be replaced by 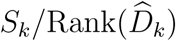. Note that the individual diagonal elements of 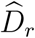 *are* uniquely defined, as the columns of *X*_*r*_ are the eigenvectors of Δ_*r*_. The scree plot obtained in this way is a quick and easy way to see which submodels explain a reasonable fraction of the variability we would expect to see in an ordination using distance Δ. Additionally, the residual distance matrix Δ_*r*_ can also be used for ordination if we wish visualize the observations after removing the (linear) effects of covariates from a distance matrix Δ.

A further connection to PERMANOVA can be made by noting that, as long as Δ is not rank-deficient (i.e., as long as *X*_*k*_ is in the range of Δ), the column vectors *X*_*k*_ span the range of 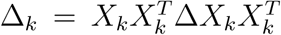 for *any* Δ. In fact, when constructing *X*, the columns *X*_*k*_ could be replaced by the eigenvectors of Δ_*k*_ having non-zero eigenvalues (denoted, say, by 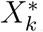); note this would not change the sums-of-squares test statistics since *X*_*k*_ and 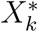 are each orthonormal and span the same space, and are hence related by an orthogonal transformation. In either case, 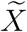 has columns that span the model matrices (the Δ_*k*_s) and the residual (Δ_*r*_) matrix of a PERMANOVA analysis. Of course, if *X*_*k*_ is comprised of a single column, then *X*_*k*_ automatically corresponds to the single eigenvector of Δ_*k*_ having non-zero eigenvalue.

Even if we choose 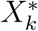 for the columns of 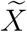, note that the diagonal elements of 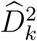 in the LDM are *not* eigenvalues of the distance matrix Δ, but rather represent the amount of variability in the OTU table *Y* that is explained by the (Euclidean) projection of the columns of *Y* on *X*_*k*_. Thus, the diagonal elements of 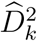 only correspond to the eigenvalues in a PERMANOVA analysis if the distance matrix Δ is the Euclidean distance. In some cases, it is possible to convert a distance matrix Δ into the Euclidean distance matrix by data transformation; in these cases, the LDM can also be considered distance-based. However, choosing 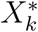 when constructing 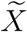 does have an advantage when constructing the scree plot, as it gives an unambiguous meaning to the individual diagonal elements of diagonal elements of 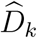 for submodels with more than one component.

### Assessing significance by permutation

We assess the significance of our test statistics, *F*_*kj*_ and *F*_*k*,global_, using a variant of a permutation scheme described by Freedman and Lane [1983]. The original Freedman-Lane procedure permutes residuals. To preserve the correlation structure among OTUs, we use the same permutation scheme for each OTU (i.e., permuting the residual matrix by row). We show in Supplementary Text S1 that this is equivalent to a permutation procedure in which the residuals are held fixed but the covariates (i.e., the rows of *X*) are permuted.

To describe our permutation approach, define *Y*_*k*_, the residual matrix obtained after fitting a reduced model to *Y* that excludes the *k*th submodel term *X*_*k*_, to be 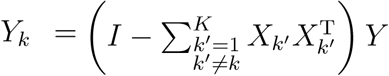 and note that, because of the orthogonality of the columns of *X*, we can write 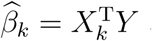 as 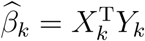. As a result, *F*_*kj*_ in (6) can be rewritten as

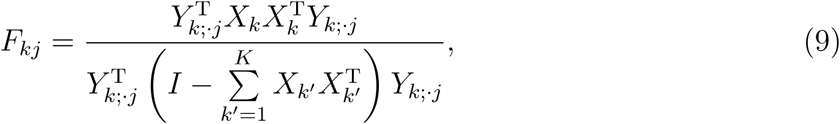

where *Y*_*k*;·*j*_ is the *j*th column of *Y*_*k*_. The analogous result for the global test statistic *F*_*k*,global_ is obtained by summing numerator and denominator over *j*. In Supplementary Text S1, we show that if *P*_*π*_ is a permutation matrix corresponding to *π*, a permutation of the integers 1, …, *n*, then the Freedman-Lane procedure is equivalent to forming the test statistics

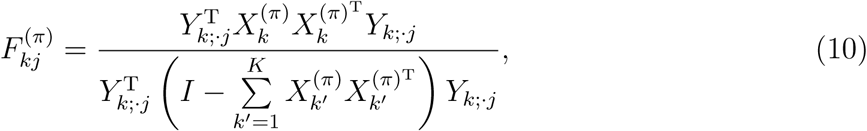

where 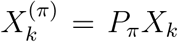 is a row-permuted version of *X*_*k*_. The test statistic for the global test 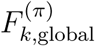 is obtained by (separately) summing the numerator and denominator of (10) over OTUs and can be written as

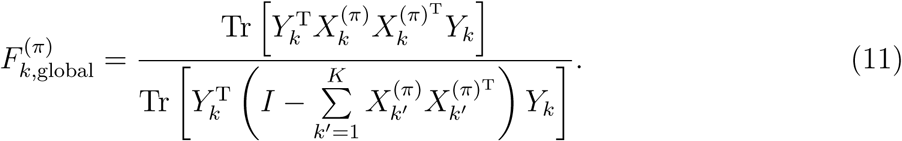

Although Freedman and Lane [1983] only considered residuals from independent observations, some simple but important cases involving correlated data can be tested using the Freedman-Lane approach. Considering permutation of covariates rather than residuals, the main requirement for a valid permutation replicate dataset is that the dataset preserve the correlation found in the original data. Thus, variables that vary within clusters (sometimes called “plots” in the Ecology literature) can be permuted within each cluster; here, we only consider the case of clustered data in which residuals within each cluster can be considered as *exchangeable*. For example, if each cluster consists of a “before treatment” observation and an “after treatment” observation from the same individual, the effect of treatment can be tested by randomly permuting the “before” and “after” assignment within each cluster (individual). Note that in this situation, the cluster sizes need not be balanced (i.e., have equal size). If *all* variables in an analysis are constant for all cluster members (i.e., are assigned at or “above” the cluster level), only permutation replicates that assign the same value to each cluster member are allowed; here exchangeability is not required. For example, in a rodent study of the effect of diet on the gut microbiome, rodents housed in the same cage should be treated as a cluster, as rodents are coprophagic. Thus, when permuting diet, rodents in the same cage should always be assigned the same diet. Note that for datasets with variables assigned at or above the cluster level, the cluster sizes must all be equal or the data must be stratified by cluster size with all permutations taking place within strata. Our implementation of the LDM uses the same permutation options available in the R package `vegan`, through the R package `permute`.

Our software implementation of the LDM uses sequential stopping rules to increase computational efficiency. When only the global test is of interest, we adopt the sequential stopping rule of Besag and Clifford [1991] for calculating the *p*-value of the global test. This algorithm terminates when either a pre-determined number *L*_min_ of rejections (i.e., the permutation statistic exceeded the observed statistic) has been reached or a pre-determined maximum number *K*_max_ of permutations have been generated. When the OTU-specific results are desired, we use the algorithm proposed by Sandve *et al.* [2011], which adds a FDR-based sequential stopping criterion to the Besag et al. algorithm. Note that the Sandve et al. algorithm limits the total number of needed permutations to *J* × *L*_min_ × *α*^−1^, which is 200 times the number of OTUs *J* when the nominal FDR *α* = 10% and *L*_min_ = 20. When testing multiple hypotheses (e.g., both the global and OTU-specific hypotheses, or hypotheses corresponding to multiple submodels), we generate permutations until all hypotheses reach their stopping point.

### A Freedman-Lane PERMANOVA test (PERMANOVA-FL)

The operations that lead to the *F*-statistics for the LDM can also be used to develop an improved PERMANOVA test statistic. We first consider a Euclidean distance Δ. Following McArdle and Anderson [2001], we note that any Euclidean distance Δ can be written as Δ = *ZZ*^T^ for some matrix *Z*; we then write a linear model of the form (3) in which *Z* replaces *Y*. Here the only tests of interest are the global tests; the analogues of the OTU-specific tests are tests of the effect of covariates on the *j*^th^ component (column) of *Z* and are only used as intermediate steps. After replacing *Y* with *Z* in (7) and using the invariance of the trace to cyclic permutations, the statistic *F*_*k*,global_ can be rewritten as

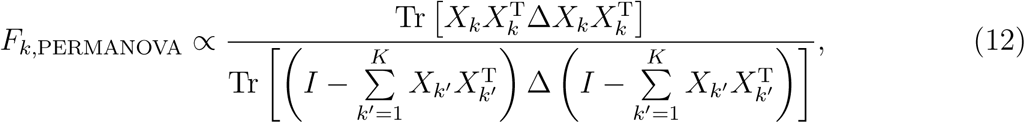

which is the usual form of the PERMANOVA *F* statistic; note that Δ here can be generalized to any (non-Euclidean) distance Δ. The same argument leading to (9) yields

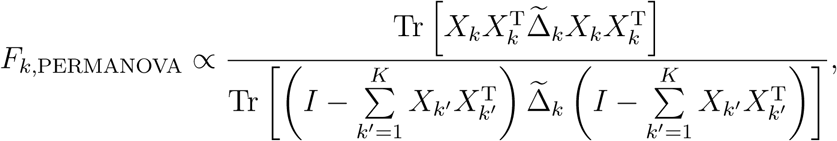

where

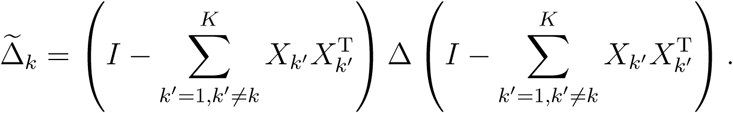

Thus, for a replicate dataset having covariates 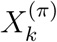, the Friedman-Lane PERMANOVA test statistic can be obtained by replacing *Y* by *Z* in (11):

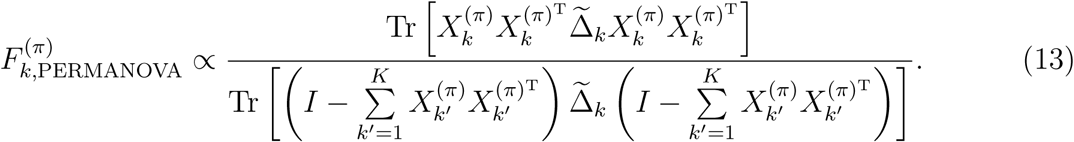

We refer to this test as PERMANOVA-FL. The same kinds of restricted permutations as in our implementation of the LDM are available in PERMANOVA-FL.

The permutation scheme implemented in `adonis` is similar to (13) except that the 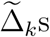 are all replaced by Δ. We further note that our proposed permutation replicates in (13) have the same advantages as the PERMANOVA replications implemented in `adonis`, in that they only require functions of the distance matrix Δ (which, in our approach, are the projected distance matrices 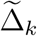). As a result, our approach, like other implementations of PERMANOVA, can be computed even if the distance matrix is non-Euclidean. Further, the distance matrices 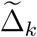 do not need to be recalculated for each replicate.

### The arcsin-root transformation

The LDM can also be applied to transformed data. Because we consider frequency data, we show we achieve good results using the arcsin-root transformation, which is variance-stabilizing for Multinomial and Dirichlet-Multinomial (DM) counts. Thus we write 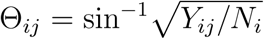 where *Y*_*ij*_ are the raw counts and *N*_*i*_ are the library sizes. We can Θ additionally center Θ, replacing it by (*I* − *n*^−1^11^T^) Θ if we also plan to center Δ. We can now replace *Y* by Θ in (1) or (3) and proceed as before. This approach is related to an approach of Berkson [1944] for fitting logistic models to bioassay data. We also had considered a logit-based model using Haldane’s [Haldane, 1956] unbiased logit by forming Θ_*ij*_ = ln{(*Y*_*ij*_ + 0.5)*/*(*N*_*i*_ − *Y*_*ij*_ + 0.5)} but found that the arcsin-root transform performed better in all cases we examined. We expect the LDM applied to (untransformed) frequency data will work best when the associated OTUs are abundant, while we expect the LDM applied to arcsin-root-transformed frequencies to work best when the associated OTUs are less abundant. Since we do not know the association mechanism *a priori*, we also consider an omnibus strategy that simultaneously applies LDM on both data scales. For the omnibus tests, we use the minimum of the *p*-value obtained from the frequency and arcsin-root-transformed data as the final test statistic and use the corresponding minima from the permuted data to simulate the null distribution [Westfall and Young, 1993]. As demonstrated by our simulation studies, the omnibus test achieves the best overall performance compared to the LDM applied to either data scale and is recommended for use in real data analysis.

## Results

### Simulation studies

We conducted several simulation studies to evaluate the performance of the LDM and compare it to competing methods. To evaluate the global test, we compared our results to those obtained using our own implementation of PERMANOVA and the PERMANOVA implemented in `adonis2`. We also calculated OTU-specific tests using the LDM, which we compared to results from DESeq2. We only performed limited comparisons to aMiSPU, edgeR, MetagenomeSeq, Wilcoxon rank-sum test (applied to OTU frequencies), ANCOM, and ALDEx2, because aMiSPU does not allow for clustered data, edgeR generally has highly inflated FDR [Hawinkel *et al.*, 2017], and the others do not allow for confounding covariates.

To generate our simulation data, we used the same motivating dataset as Zhao *et al.* [2015], specifically data on the upper-respiratory-tract (URT) microbiome first described by Charlson *et al.* [2010]. To simulate read count data for the 856 OTUs reported in this study, we adopted a DM model using the empirical frequencies calculated from the study data; we set the overdispersion parameter to the estimate 0.02 obtained from these data, which is also the median value we observed in an admittedly brief survey of the literature [Chen and Li, 2013, La Rosa *et al.*, 2012, Morgan *et al.*, 2015]. While the original microbiome dataset was generated from 454 pyrosequencing with mean library size ∼1500, we increased the mean library size to 10000 to reflect Illumina MiSeq sequencing which is currently in common usage. For each simulation, we generated data for 100 samples unless otherwise noted. We also conducted sensitivity analysis with a wide range of library sizes, overdispersion parameters, and sample sizes, and by replacing the DM model with a Poisson log-normal model (PLNM) or Negative-Binomial (NB) model to generate the read count data (the PLNM and NB model are described in Supplementary Texts S2 and S3, respectively).

We focused on two complementary scenarios. The first scenario (S1) assumed that a large number of moderately abundant and rare OTUs were differentially abundant between cases and controls, and the second scenario (S2) assumed the top 10 most abundant OTUs were differentially abundant. Both scenarios have a one-way, case-control design with a confounder and independent samples. Later we varied these scenarios to simulate a continuous trait, a two-way design, or clustered data.

In both scenarios S1 and S2, we let *U* denote case-control status and assumed an equal number of cases (*U* = 1) and controls (*U* = 0). We simulated a confounder, *C* = 0.5*U* + *ϵ*, where *ϵ* was drawn from a uniform distribution on [0, 1]. In S1, we uniformly and independently sampled two (overlapping) sets of 428 OTUs (half of all OTUs), the first set associated with *U* and the second set associated with *C*; the set for *U* was sampled after excluding the top three most abundant OTUs to focus on less abundant OTUs. In S2, we assumed the ten most abundant OTUs were associated with *U* and the next forty most abundant OTUs were associated with *C*. These OTU sets were held fixed across replicates of data. We denoted the OTU frequencies estimated from the real data by the vector *π*_1_ and formed vectors *π*_2_ and *π*_3_ by first setting *π*_2_ and *π*_3_ equal to *π*_1_ and then randomly permuting those frequencies in *π*_2_ and *π*_3_ that belong to the selected set of OTUs associated with *U* and *C*, respectively. Note that the frequencies for OTUs not selected to be associated with *U* (or *C*) remain the same in *π*_1_ and *π*_2_ (or *π*_3_). We then defined a sample-specific frequency vector as 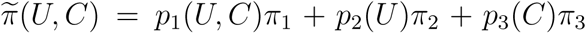, where *p*_2_(*U*) = *βU, p*_3_(*C*) = *β*_*C*_*C, p*_1_(*U, C*) = 1 − *p*_2_(*U*) − *p*_3_(*C*). In this model, *β* and *β*_*C*_ are the effect sizes of *U* and *C* on the overall community compositions, respectively; here we set *β*_*C*_ to 0.3 except for simulations with no confounding, for which we set *β*_*C*_ to zero. At the OTU level, the strengths and directions of the effects of *U* (or *C*) are heterogeneous because the resulting frequencies at each OTU are characterized not just by *β* (or *β*_*C*_), but also by the differences between *π*_1_ and *π*_2_ (or *π*_1_ and *π*_3_), which vary in magnitude and sign at different OTUs. We then generated the OTU count data for each sample using the DM model with 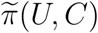, overdispersion parameter of 0.02, and library size sampled from *N*(10000, 10000*/*3) and left-truncated at 500. By mixing *π*_1_, *π*_2_, and *π*_3_ in a way that depends on the values of *U* and *C*, we induced associations between the selected OTUs and *U* and *C*. Note that *π*_1_ serves as the “reference” OTU frequencies that characterizes samples for which *U* and *C* are both zero. In addition, the correlations among *U, C*, and OTU frequencies establish *C* as a confounder of the association between *U* and the OTUs. Finally, note that when 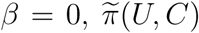 does not depend on *U*, so that the null hypothesis of no association between *U* and OTU frequencies holds. The simulations of clustered data, data with a two-way design, and data with a quantitative trait are described in Supplementary Text S4.

We evaluated type I error and power for testing the global hypothesis at nominal significance level 0.05, and we assessed empirical sensitivity (proportion of truly associated OTUs that are detected) and empirical FDR for testing individual OTUs at nominal FDR 10%. Results for type I error were based on 10000 replicates; all other results are based on 1000. In all simulations with confounders, we treated *C* and *U* as separate submodels, when fitting the LDM. For the two-way simulations, *U*_1_ and *U*_2_ were considered as separate submodels.

#### Results for testing global hypotheses with independent samples in the one-way case-control design

For testing the global hypothesis *H*_0_: *β* = 0 of no association between microbiome composition and *U*, we applied the LDM on the frequency and arcsin-root scales and also calculated the omnibus test; these results are presented as VE-freq, VE-arcsin, and VE-omni, respectively, where VE denotes variance explained. We also applied our own implementation of PERMANOVA as well as the `adonis2` implementation; we refer to them as PERMANOVA-FL and adonis2, respectively.

Table 1 (top panel) shows our results for type I error. All methods, after adjusting for confounders, had correct type I error; with the small sample size 20, the type I error rates of PERMANOVA-FL and LDM methods were slightly conservative, which is consistent with the findings of Anderson and Legendre [1999]. There was substantial inflation of type I error for S1 and modest inflation for S2 when the confounder was not accounted for, demonstrating that our methods are effective in accounting for confounders, with either modest or substantial confounding. The type I error rates were also close to 0.05 when the PLNM was adopted for count data simulation (Table S1).

**Table 1.** Type I error for testing the global hypothesis at nominal level 0.05.

| | Scenario | $n$ | PERMANOVA-FL | adonis2 | VE-freq | VE-arcsin | VE-omni |
| --- | --- | --- | --- | --- | --- | --- | --- |
| Independent samples, one-way case-control design |  |  |  |  |  |  |  |
| Adjusting for confounder | S1 | 20 | 0.040 | 0.052 | 0.045 | 0.031 | 0.038 |
|  |  | 100 | 0.053 | 0.052 | 0.050 | 0.054 | 0.053 |
|  | S2 | 20 | 0.039 | 0.054 | 0.043 | 0.026 | 0.034 |
|  |  | 100 | 0.050 | 0.050 | 0.052 | 0.047 | 0.053 |
| Not adjusting for confounder | S1 | 20 | 0.299 | 0.296 | 0.215 | 0.408 | 0.362 |
|  |  | 100 | 0.987 | 0.987 | 0.913 | 0.998 | 0.997 |
|  | S2 | 20 | 0.065 | 0.064 | 0.056 | 0.076 | 0.067 |
|  |  | 100 | 0.151 | 0.151 | 0.083 | 0.245 | 0.194 |
| Independent samples, two-way design |  |  |  |  |  |  |  |
| Testing for $U_1$ | S1 | 100 | 0.046 | 0.013 | 0.048 | 0.048 | 0.050 |
|  | S2 | 100 | 0.049 | 0.048 | 0.051 | 0.051 | 0.050 |
| Testing for $U_2$ | S1 | 100 | 0.049 | 0.036 | 0.049 | 0.046 | 0.049 |
|  | S2 | 100 | 0.053 | 0.048 | 0.053 | 0.054 | 0.054 |
| Clustered samples, one-way case-control design |  |  |  |  |  |  |  |
| Accounting for clustering | S1 | 100 | 0.050 | 0.050 | 0.048 | 0.051 | 0.053 |
|  | S2 | 100 | 0.044 | 0.047 | 0.047 | 0.044 | 0.045 |
| Not accounting for clustering | S1 | 100 | 1 | 1 | 1 | 1 | 1 |
|  | S2 | 100 | 1 | 1 | 0.999 | 1 | 1 |
| Independent samples, continuous trait |  |  |  |  |  |  |  |
| Adjusting for confounder |  | 100 | 0.049 | 0.051 | 0.055 | 0.049 | 0.051 |
| Not adjusting for confounder |  | 100 | 0.095 | 0.095 | 0.090 | 0.093 | 0.091 |

Results for PERMANOVA-FL and adonis2 are based on the Bray-Curtis distance. $n$ is the number of samples. When testing $U_1$ , we set $\beta_2 = 0.5$ ; when testing $U_2$ , we set $\beta_1 = 0.5$ . All analyses of clustered data adjust for the confounder.

Figure 1 (top panel) displays our results for power. We can see that VE-arcsin is more powerful than VE-freq under S1 and vice versa under S2; this is presumably because the variance stabilization of the arcsin-root transformation gives greater power to detect association with the rare OTUs that carry association in S1, while the untransformed data gives increased power to detect the common-OTU associations that characterize S2. In both cases, VE-omni achieved almost the same power as the most powerful test, without having to know whether common or rare OTUs were most important. PERMANOVA-FL has varying power depending on the choice of distance measure: the power is lowest with the weighted-UniFrac distance, since the association was induced without reference to any phylogenetic tree, and the power is highest with the Hellinger distance in S1 and Bray-Curtis in S2. In both S1 and S2, our best-performing method has comparable power as the best-performing PERMANOVA-FL. aMiSPU has a comparable power to VE-omni under S1 but has the least power under S2. Our sensitivity analysis showed that the relative performance of these methods persist for a wide range of library sizes, overdispersion parameters, and sample sizes (Figure S1), as well as with the PLNM (Figure S2). For this set of studies, PERMANOVA-FL and adonis2 yielded very similar power.

**Fig. 1.**
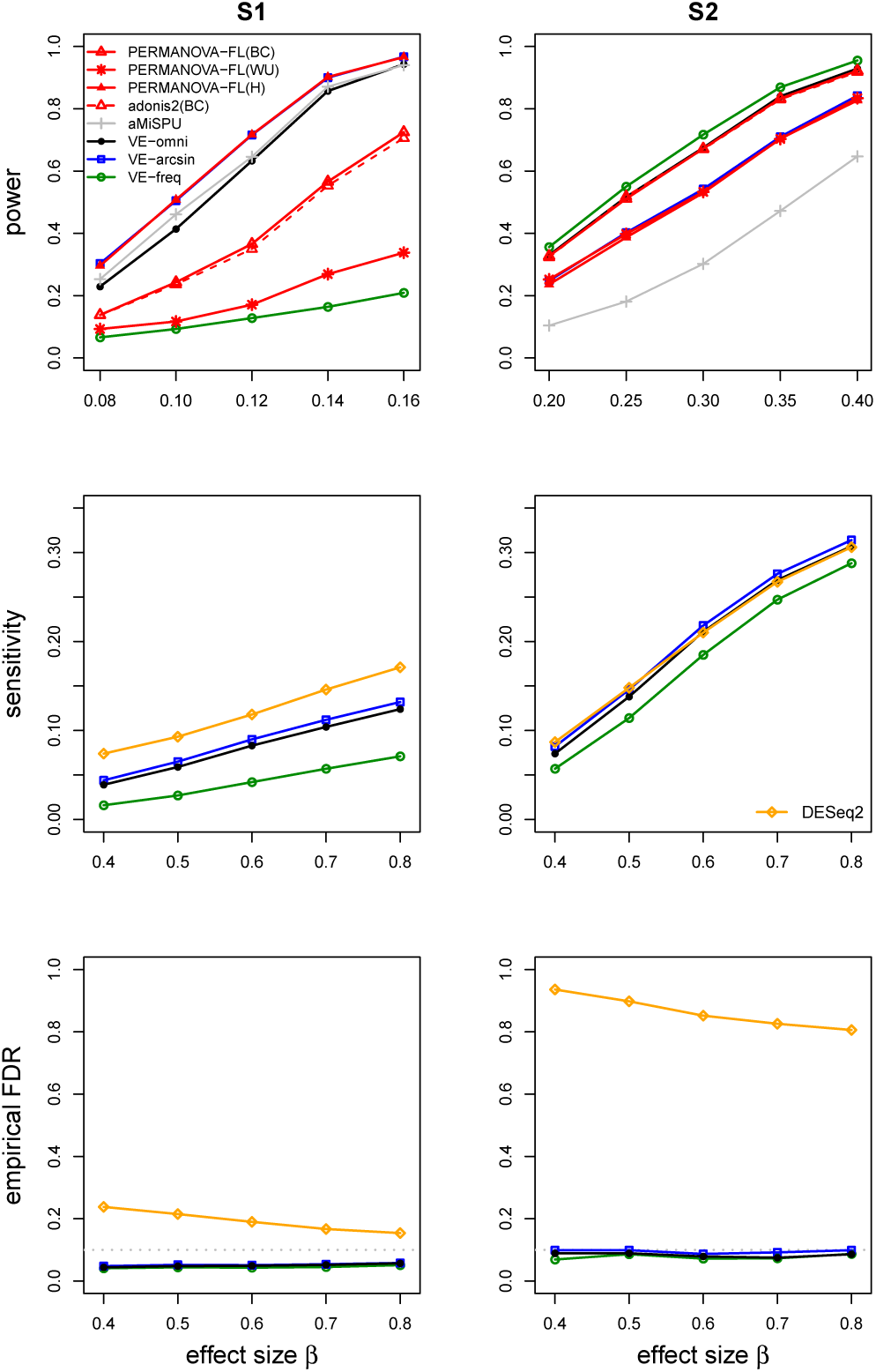
Simulation results for one-way, case-control studies with independent samples. The gray dotted lines represent FDR = 0.1. BC: Bray-Curtis; WU: weighted UniFrac; H: Hellinger.

#### Results for testing individual OTUs with independent samples in the one-way case-control design

Because PERMANOVA-FL and adonis2 does not provide OTU-specific results, we compared our results on testing individual OTUs to DESeq2. When applying DESeq2, we replaced the default normalization by GMPR normalization [Chen and Chen, 2017], which was specifically developed for zero-inflated microbiome data.

Figure 1 middle and bottom panels display results on empirical sensitivity and empirical FDR, respectively. The LDM-based methods controlled FDR at 10% in all cases; their empirical FDRs are conservative (and the sensitivity values are low) in S1 because this scenario permuted frequencies among 428 OTUs selected for *U*, majority of which are rare, and thus generated many weakly associated OTUs that are essentially null OTUs. The sensitivity of VE-omni tracks the method between VE-freq and VE-arcsin that performs better. Note that VE-arcsin has a higher sensitivity than VE-freq in both S1 and S2, but the order can be reversed in the scenario with a continuous trait (Figure S6). In contrast, the empirical FDRs for DESeq2 are modestly inflated under S1 and highly inflated under S2. Even when the read count data were simulated from the NB model (for which DESeq2 was designed), the empirical FDRs of DESeq2 are still inflated (Figure S3), likely due to the large number of zero counts.

Because MetagenomeSeq, the Wilcoxon rank-sum test, ANCOM, and ALDEx2 do not allow for adjustment of confounding covariates, we have not included results from these methods in Figure 1. To compare with these methods, we set *β*_*C*_ = 0 for both scenarios S1 and S2 to remove confounders and displayed the results in Figure S4. MetagenomeSeq always controlled FDR but was extremely conservative (FDR *<* 2% for nominal FDR of 10% and sample size 100) in detecting associated OTUs for the simulations we conducted. The Wilcoxon test controlled FDR and achieved good sensitivity when the DM model was used for generating the read count data; when the PLNM was used (with no confounders), the data appeared less overdispersed and Wilcoxon had consistently lower sensitivity than VE-omni (Figure S2). ANCOM cannot control FDR under S2. While ALDEx2 controlled FDR under S1 and S2, its sensitivity is consistently lower than the LDM-based methods. We have also included edgeR in this study, whose highly inflated FDR corroborated the finding in Hawinkel *et al.* [2017]. Finally, DESeq2 failed to control FDR even in the absence of any confounders.

#### Results for the two-way design

We set *β*_1_ = 0 and *β*_2_ = 0.5 to ascertain the type I error of the test of *U*_1_, and *β*_2_ = 0 and *β*_1_ = 0.5 to ascertain the type I error of the test of *U*_2_, which results were summarized in Table 1. Both the LDM and PERMANOVA-FL yielded correct type I error for testing each factor, whereas adonis2 had conservative type I error in scenario S1. The type I error using adonis2 was about a factor of 3 smaller for testing *U*_1_ than for testing *U*_2_ because the sampled OTUs for association with *U*_2_ (or *C*) included the top two most abundant OTUs and, as a result, *U*_2_ had a stronger global effect on the OTUs than *U*_1_. Consistent with the conservative type I error, adonis2 had lower power than PERMANOVA-FL (Figure 2). LDM (VE-omni) continued to maintain good power relative to PERMANOVA-FL for either factor (Figure 2). Further, LDM controlled FDR for OTUs that were detected to be associated with either factor (Figure 2).

**Fig. 2.**
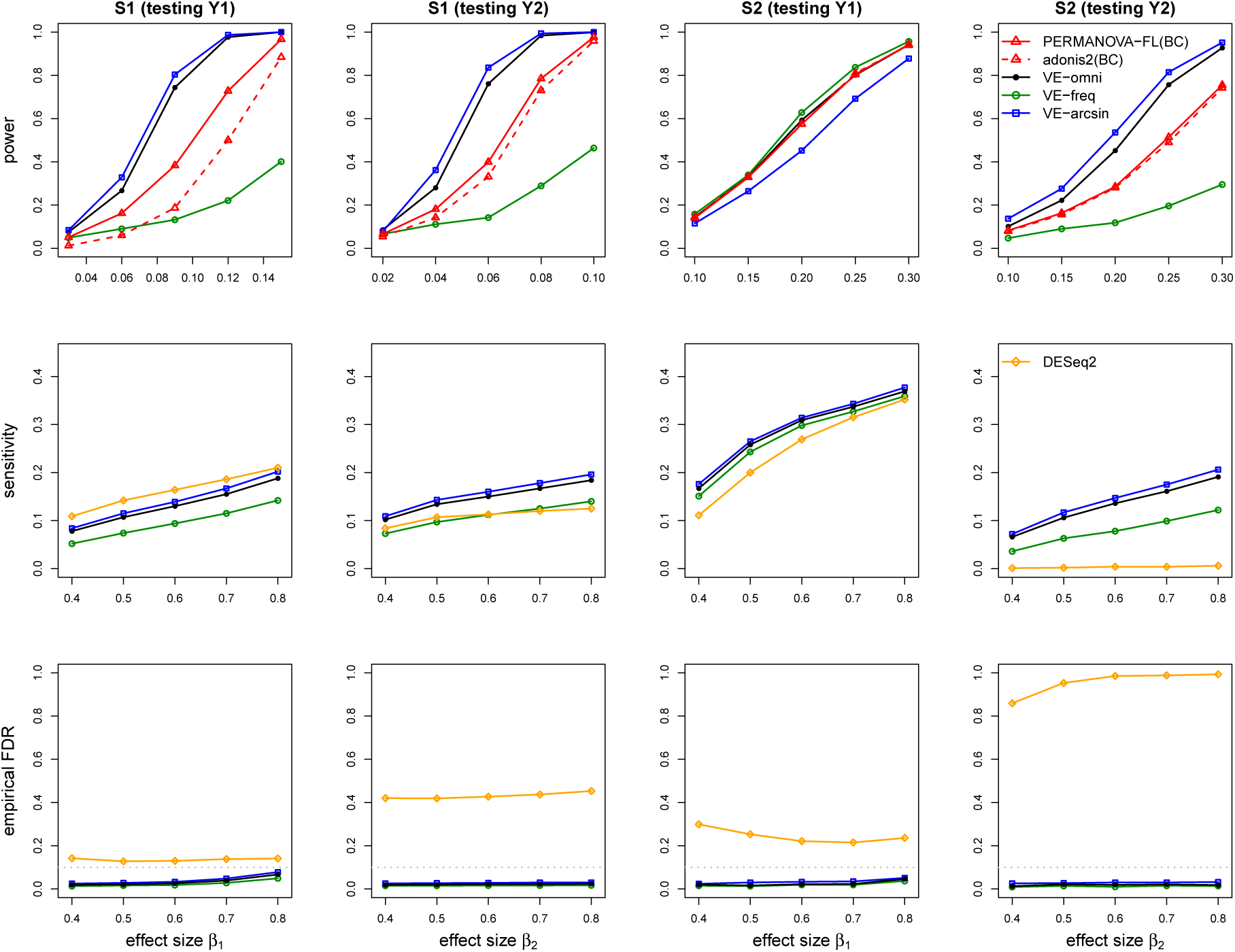
Simulation results for studies with the two-way design and independent samples. BC: Bray-Curtis. When testing *U*_1_, we set *β*_2_ = 0.5; when testing *U*_2_, we set *β*_1_ = 0.5.

#### Results for clustered data

In Table 1, we can see that permuting the case-control status over clusters rather than observations yields the correct type I error for all methods. We also calculated the type I error we would have obtained if we had incorrectly ignored the clustering structure when performing the permutations. Note that failure to account for the clustering structure result in a type I error of 100%. In Figure S5, we see the LDM controlled FDR for these data, although the power and sensitivity is lower than was observed with the same number of samples which were unclustered (Figure 1). This is reasonable, as data with within-individual correlation is typically not as informative as data from an equivalent number of independent samples.

#### Results for continuous trait

From Table 1, we again see that all methods (adjusting for the confounder) have the correct type I error for data with a continuous trait; there was inflation of type I error when the confounder was not accounted for. In Figure S6, we see that the power of most methods is about the same. Although the sensitivity remains low as the effect size *β* increases, this appears to be related to the sample size, as we also show that the sensitivity increases rapidly as the sample size increases (at fixed *β* = 3). The LDM continues to control FDR as the sample size and sensitivity increase, while the empirical FDR for DESeq2 is never less than 40% for the range of sample sizes we considered.

### Analysis of two microbiome datasets

To show the performance of the LDM in real microbiome data, we reanalyzed two datasets that were previously analyzed using MiRKAT and MMiRKAT [Zhan *et al.*, 2017] (a variant of MiRKAT for testing association between multiple continuous covariates and microbiome composition). The first is from a study of the association between the upper-respiratory-tract (URT) microbiome and smoking, and the second is from a study of the association between the prepouch-ileum (PPI) microbiome and host gene expression in patients with inflammatory bowel disease (IBD). We compared the performance of our global test (VE-omni) with PERMANOVA-FL, MiRKAT, and MMiRKAT; we also compared our OTU-specific results with results from DESeq2.

#### URT microbiome and smoking association study

The data for our first example were generated as part of a study to examine the effect of cigarette smoking on the orpharyngeal and nospharyngeal microbiome [Charlson *et al.*, 2010]. The 16S sequence data are summarized in an OTU table consisting of data from 60 samples and 856 OTUs with mean library size 1500; metadata on smoking status (28 smokers and 32 nonsmokers) and two additional covariates (gender and antibiotic use within the last 3 months) was also available. An imbalance in the proportion of male subjects by smoking status (75% in smokers, 56% in non-smokers) indicates potential for confounding. Zhao *et al.* [2015] analyzed these data using MiRKAT, finding a significant global association between microbiome composition and smoking status after adjusting for gender and antibiotic use. We used the Bray-Curtis distance for our analysis because it led to the smallest *p*-values compared to other distances in Zhao et al. (2015). We combined gender and antibiotic use into a single submodel and treated smoking status as another submodel when fitting the LDM.

We first constructed the ordination plots in Figure 3 using the Bray-Curtis distance after removing the effects of gender and antibiotic use (i.e., using Δ_1_ as the distance matrix); these plots demonstrate a clear shift in smokers compared with nonsmokers even after removing the effect of potential confounders. The accompanying scree plots (Figure 3) on both frequency and arcsin-root scales further suggest that smoking explains an important faction of the variability in the OTU table. The residual (non-model) components are plotted in decreasing order of the size of the eigenvalue of the component in the spectral decomposition of Δ_2_ (after removing the effect of confounders and smoking); the high correlation between the order of values *D*_*k*_ from the LDM and the order of eigenvalues of Δ_2_ is noteworthy. We filtered out OTUs with presence in less than 5 samples, retaining 233 OTUs for analysis. The results of the LDM global tests, along with results from PERMANOVA-FL and MiRKAT, are presented in top-left panel of Table 2. VE-omni gave a smaller *p*-value than MiRKAT or PERMANOVA-FL based on the Bray-Curtis distance. In the top-right panel of Table 2, we show the results of our OTU-specific tests. VE-omni detected 5 OTUs (which include the 4 OTUs detected by VE-freq and constitute 5 of the 14 OTUs detected by VE-arcsin) whereas DESeq2 detected none. The inefficiency of DESeq2 is consistent with our simulation studies when the mean library size was 1500 (results not shown).

**Table 2.**
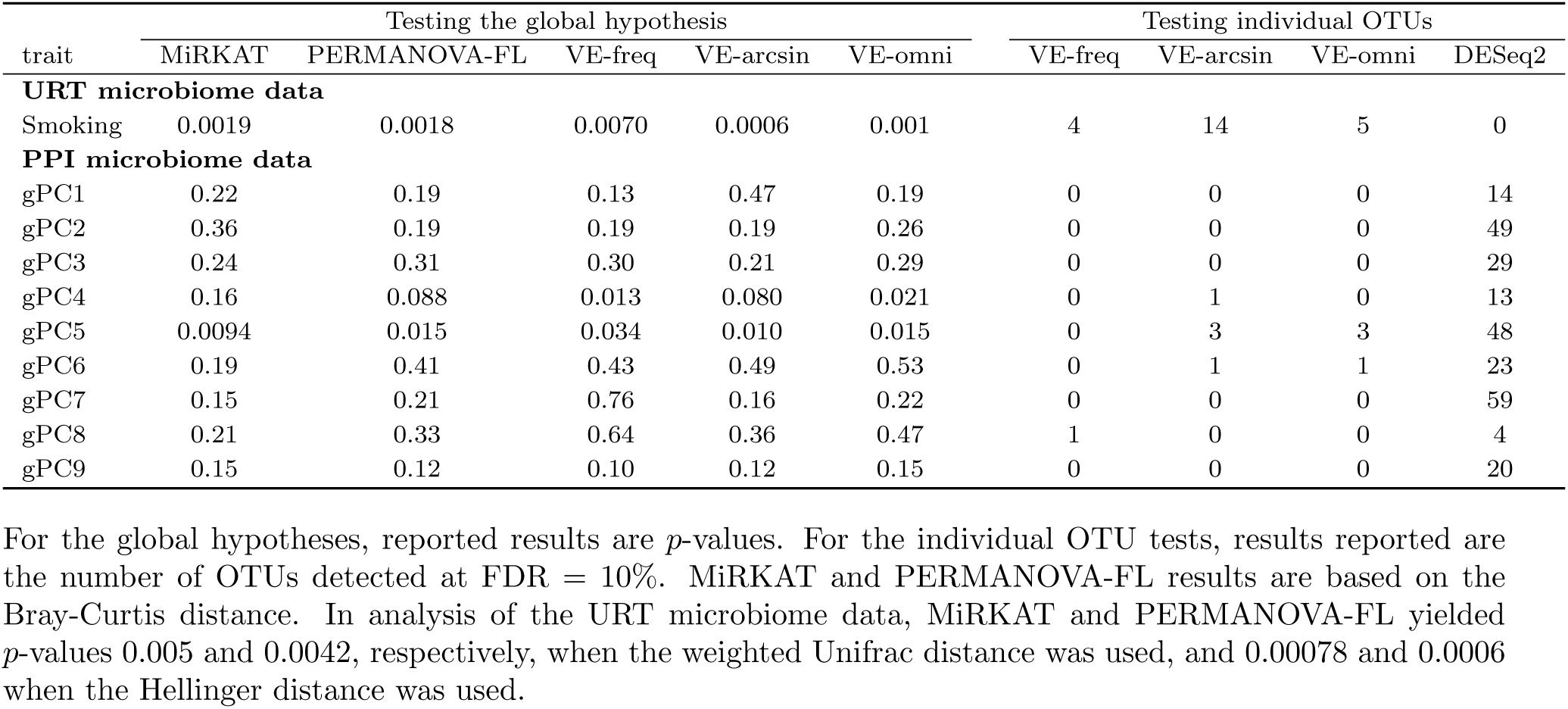
Results in analysis of the two real datasets.

| trait | Testing the global hypothesis |  |  |  |  | Testing individual OTUs |  |  |  |
| --- | --- | --- | --- | --- | --- | --- | --- | --- | --- |
|  | MiRKAT | PERMANOVA-FL | VE-freq | VE-arcsin | VE-omni | VE-freq | VE-arcsin | VE-omni | DESeq2 |
| **URT microbiome data** |  |  |  |  |  |  |  |  |  |
| Smoking | 0.0019 | 0.0018 | 0.0070 | 0.0006 | 0.001 | 4 | 14 | 5 | 0 |
| **PPI microbiome data** |  |  |  |  |  |  |  |  |  |
| gPC1 | 0.22 | 0.19 | 0.13 | 0.47 | 0.19 | 0 | 0 | 0 | 14 |
| gPC2 | 0.36 | 0.19 | 0.19 | 0.19 | 0.26 | 0 | 0 | 0 | 49 |
| gPC3 | 0.24 | 0.31 | 0.30 | 0.21 | 0.29 | 0 | 0 | 0 | 29 |
| gPC4 | 0.16 | 0.088 | 0.013 | 0.080 | 0.021 | 0 | 1 | 0 | 13 |
| gPC5 | 0.0094 | 0.015 | 0.034 | 0.010 | 0.015 | 0 | 3 | 3 | 48 |
| gPC6 | 0.19 | 0.41 | 0.43 | 0.49 | 0.53 | 0 | 1 | 1 | 23 |
| gPC7 | 0.15 | 0.21 | 0.76 | 0.16 | 0.22 | 0 | 0 | 0 | 59 |
| gPC8 | 0.21 | 0.33 | 0.64 | 0.36 | 0.47 | 1 | 0 | 0 | 4 |
| gPC9 | 0.15 | 0.12 | 0.10 | 0.12 | 0.15 | 0 | 0 | 0 | 20 |

For the global hypotheses, reported results are $p$ -values. For the individual OTU tests, results reported are the number of OTUs detected at $\text{FDR} = 10\%$ . MiRKAT and PERMANOVA-FL results are based on the Bray-Curtis distance. In analysis of the URT microbiome data, MiRKAT and PERMANOVA-FL yielded $p$ -values 0.005 and 0.0042, respectively, when the weighted Unifrac distance was used, and 0.00078 and 0.0006 when the Hellinger distance was used.

**Fig. 3.**
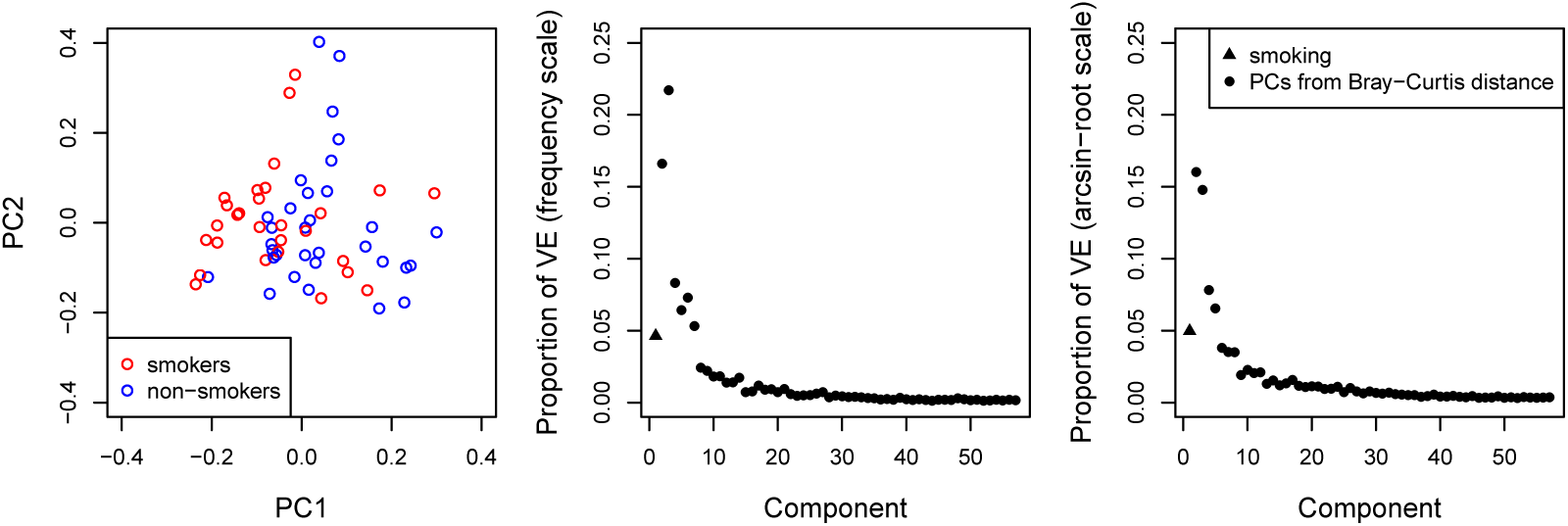
Exploratory analysis of the URT microbiome data based on the Bray-Curtis distance. Left plot: ordination plot after removing the effects of confounders gender and antibiotic use, colored by smoking status. Center and right plots: proportions of variance explained by smoking and the PCs of the (residual) distance measure after removing the effects of gender and antibiotic use; the PCs are ordered by their Bray-Curtis eigenvalues. The center plot is based on frequency data and the right plot is based on arcsin-root transformed data. The components are ordered by the magnitude of their corresponding eigenvalue in a spectral decomposition of Δ_2_ (the distance matrix after removing the effect of the confounders and smoking).

#### PPI microbiome and host gene expression association study

The data for our second example were generated in a study of the association between the mucosal microbiome in the prepouch-ileum (PPI) and host gene expression among patients with IBD [Morgan *et al.*, 2015]. The PPI microbiome data are summarized in an OTU table with data from 196 IBD patients and 7,000 OTUs; gene expression data at 33,297 host transcripts, as well as clinical metadata such as antibiotic use (yes/no), inflammation score (0–13), and disease type (familial adenomatous polyposis/FAP and non-FAP) were also available. The data also included nine gene principal components (gPCs) that together explain 50% of the total variance in host gene expression. Zhan *et al.* [2017] gave a joint test of all nine gPCs for association with microbiome composition, using MMiRKAT based on the Bray-Curtis distance measure and adjusting for antibiotic use, inflammation score, and disease type (FAP/non-FAP). Here we performed the same joint test using the LDM by putting the confounders in one submodel and then including all nine gPCs in another submodel; however, we followed Morgan *et al.* [2015] in only analyzing the original 196 PPI samples, not an additional 59 pouch samples from some of the same individuals included in the analysis of Zhan *et al.* [2017]. We filtered out OTUs found in fewer than 5% of samples, retaining 2096 OTUs for analysis. VE-freq, VE-arcsin, and VE-omni yielded *p*-values 0.023, 0.0084, and 0.015, respectively, and detected 0, 4, and 3 OTUs (the 3 OTUs detected by VE-omni are included in the 4 OTUs detected by VE-arcsin) that significantly accounted for the global association at a nominal FDR rate of 10%. PERMANOVA-FL had *p*-value 0.0076 and MMiRKAT 0.0049, both based on the Bray-Curtis distance.

We also followed Zhan et al. (2016) to conduct individual tests of each of the nine gPCs. We treated each gPCs as a separate submodel in a single LDM, with the first submodel accounting for the same confounders as in the joint test. Note that the gPCs are orthogonal. Scree plots for frequency and arcsin-root transformed data are shown in Figure S7, and indicate that gPC4 and gPC5 are most likely to be associated with microbial composition although any association is likely to be marginal. In fact, Table 2 confirms that gPC4 and gPC5 showed significant associations (at the 0.05 significance level) with the overall microbiome composition by the global tests; no other gPCs were found to be associated. Both VE-omni and VE-arcsin detected (the same) 3 associated OTUs for gPC5, while VE-arcsin additionally found one OTU associated with gPC4. Both VE-omni and VE-arcsin also detected (the same) 1 OTU for gPC6, which was not significantly associated with the microbiome in a global test by any method. In contrast to the results obtained by the LDM, DESeq2 detected between 4 and 59 OTUs for each of the nine gPCs, which seems implausible given the results of the global tests. These findings may be related to the failure of DESeq2 to control FDR in the presence of confounders in our simulation studies.

## Discussion

We have presented the LDM, a linear model for testing association hypotheses for microbiome data that can account for the complex designs found in microbiome studies. We have shown that the LDM has good power for global tests of association between variables of interest and the microbiome when compared to existing methods such as PERMANOVA and MiRKAT (the simulation results of MiRKAT were similar to those of PERMANOVA and thus not shown), but also provides OTU-specific tests. This is true even when confounding covariates are present, or when the study design results in correlated data. We have additionally shown that the OTUs identified by the LDM preserve FDR, while those identified by RNA-Seq-based approaches such as DESeq2 typically do not; further, since global and OTU-specific tests are unified, our analysis of the PPI microbiome data show that the LDM is less likely to identify “significant” OTUs for variables that are not globally significant. In the analyses we show here, there was only one instance where the LDM discovered OTUs that were significantly associated with a variable but the LDM global test for that variable was non-significant (gPC6 in the PPI data); in our simulations there were no such cases, although that may be because we only evaluated sensitivity for effect sizes that were large enough that the global test was always positive. While some analysts may choose to only calculate OTU-specific tests in the presence of a significant global test, this restriction is not required to control FDR at the OTU level. We have evaluated our approach using simulated data with realistic amounts of overdispersion, confounding covariates and clustered data, and have shown how it can be applied to two real datasets.

We have implemented our approach in the R package LDM, which is scalable to large sample sizes. Using a single thread of a MacBook Pro laptop (2.9 GHz Intel Core i7, 16 GB memory) and the default value *L*_min_ = 20, it took 8, 19, 833 seconds to perform integrated global and OTU tests with a simulated dataset that consists of 20, 100, and 1000 samples. In our applications to real data, we used *L*_min_ = 100 to ensure stability of results which are based on Monte Carlo sampling. It took 23 seconds (and stopped after 29400 permutations) to perform global and OTU tests with the URT data, and 5 hours (and stopped after 1273000 permutations) to perform global and OTU tests of nine gPCs separately with the PPI data. Because the LDM has dual goals of testing both the global and OTU-specific hypotheses, and because it is based on permutation, it generally takes more run time than tests that only test the global hypotheses, such as PERMANOVA, or OTU-specific tests that are based on analytical *p*-values, such as DESeq2. However, because our R package LDM uses a sequential stopping rule, the computational time is still manageable. In particular, LDM is much faster than ANCOM and comparable to ALDEx2.

The LDM has features in common with RA (Redundancy Analysis), a multivariate technique to describe how much variability in one matrix (say, an OTU table *Y*) can be described by variables in a second matrix (say, a design matrix *X*); see more details in Supplementary Text S5. The LDM differs from RA most importantly in its ability to simultaneously obtain results for several submodels. To fit more than one submodel using RA, it is necessary to fit RA to each submodel, using data for which the previous submodels have been projected off. This precludes use of type III (last variable added) tests, which are known to be the most powerful [Muller and Fetterman, 2012]. Our use of the Freedman-Lane approach also gives superior performance; in simulations, our PERMANOVA-FL had higher power than the `adonis` and `adonis2` functions in the *R* package `vegan`, even though `adonis2` is based on some form of permutation of residuals (according to `adonis2` output).

Although the LDM is primarily based on the Euclidean distance in its focus on sums of squares, variability explained and *F*-like tests, we have shown how information on an arbitrary distance can be incorporated in exploratory analyses, and how a distance matrix can be used to choose analysis directions when submodels contain multiple components. Although the Euclidean distance has been criticized when used for ecological analysis, Chao and Chiu [2016] have recently suggested the problems associated with use of the Euclidean distance as a measure of beta diversity are related to normalization, rather than any intrinsic failure of the Euclidean distance. Finally, while *distance-based* Redundancy Analysis [Legendre and Anderson, 1999] does incorporate distance information, like PERMANOVA, it removes information on the effects of individual OTUs, and so was not included in our discussion.

In our examples here, we have put all confounders into the first submodel *X*_1_. This conforms with practice in epidemiology in which confounders are not tested for inclusion into a model, but rather are included based on subject-area knowledge [VanderWeele and Shpitser, 2011]. With this in mind, our implementation of the LDM does not provide *p*-values for the set of variables that are designated as confounders, which makes the code run faster. However, for those who want to estimate and test the individual effects of confounders, each confounder can be treated as separate submodel, and the LDM will calculate a *p*-value for each confounder. The results obtained in this way for the remaining variables are identical.

Among OTU-specific tests in absence of confounders, we found that metagenomeSeq controlled FDR in the simulations we conducted, while Hawinkel *et al.* [2017] claimed that metagenomeSeq failed to control FDR. We noticed that Hawinkel *et al.* [2017] adopted the zero-inflated Gaussian mixture distribution (i.e., the `fitZig` function), whereas we adopted the zero-inflated log-normal mixture model (i.e., the `fitFeatureModel` function) as recommended by the metagenomeSeq R package. We also found that the Wilcoxon rank-sum test is a robust and powerful choice for detecting differentially abundant OTUs when testing a single binary covariate. A recently-developed version of the rank-sum test [Satten *et al.*, 2018] that uses inverse-probability-of-treatment weights could provide an interesting extension for categorical testing when adjustment for confounding covariates is required. However, OTU-specific tests based on the rank-sum test do not provide coherent results with any global test.

The question of whether rare OTUs should be removed before analyzing microbiome data is unresolved. In our simulations, we did not filter out any OTUs to show the validity of the LDM even in the most challenging situation where these rare OTUs are retained. We applied filters in the real data analysis to reduce the multiple-comparison penalty and increase the chance of finding OTU-level associations. Retaining only those OTUs found in at least five samples, or in at least 5% of samples, are both commonly used filters in the literature, and we used one for each data analysis. Filtering out the very rare OTUs will only minimally affect the performance of our global test because these OTUs explain a negligible proportion of the variability in either the frequency-scale or arcsin-root-transformed data.

## Disclaimer

The findings and conclusions in this report are those of the authors and do not necessarily represent the official position of the Centers for Disease Control and Prevention.

## References

Aitchison, J. (1986). The statistical analysis of compositional data. Chapman and Hall, London-New York.

Anderson, M. J. (2001). A new method for non-parametric multivariate analysis of variance. Austral ecology, 26(1), 32–46.

Anderson, M. J. and Legendre, P. (1999). An empirical comparison of permutation methods for tests of partial regression coefficients in a linear model. Journal of statistical computation and simulation, 62(3), 271–303.

Berkson, J. (1944). Application of the logistic function to bio-assay. Journal of the American Statistical Association, 39(227), 357–365.

Besag, J. and Clifford, P. (1991). Sequential Monte Carlo p-values. Biometrika, 78(2), 301–304.

Chao, A. and Chiu, C.-H. (2016). Bridging the variance and diversity decomposition approaches to beta diversity vis similarity and differentiation measures. Methods in Ecology and Evolution, 7(8), 919–928.

Charlson, E. S., Chen, J., Custers-Allen, R., Bittinger, K., Li, H., Sinha, R., Hwang, J., Bushman, F. D., and Collman, R. G. (2010). Disordered microbial communities in the upper respiratory tract of cigarette smokers. PloS one, 5(12), e15216.

Chen, E. Z. and Li, H. (2016). A two-part mixed-effects model for analyzing longitudinal microbiome compositional data. Bioinformatics, 32(17), 2611–2617.

Chen, J. and Chen, L. (2017). Gmpr: A novel normalization method for microbiome sequencing data. bioRxiv, page 112565.

Chen, J. and Li, H. (2013). Variable selection for sparse dirichlet-multinomial regression with an application to microbiome data analysis. The annals of applied statistics, 7(1).

Fernandes, A. D., Reid, J. N., Macklaim, J. M., McMurrough, T. A., Edgell, D. R., and Gloor, G. B. (2014). Unifying the analysis of high-throughput sequencing datasets: characterizing rna-seq, 16s rrna gene sequencing and selective growth experiments by compositional data analysis. Microbiome, 2(1), 15.

Freedman, D. and Lane, D. (1983). A nonstochastic interpretation of reported significance levels. Journal of Business & Economic Statistics, 1(4), 292–298.

Gower, J. C. (1966). Some distance properties of latent root and vector methods used in multivariate analysis. Biometrika, 53(3-4), 325–338.

Haldane, J. (1956). The estimation and significance of the logarithm of a ratio of frequencies. Annals of human genetics, 20(4), 309–311.

Hawinkel, S., Mattiello, F., Bijnens, L., and Thas, O. (2017). A broken promise: microbiome differential abundance methods do not control the false discovery rate. Briefings in bioinformatics.

Hu, J., Koh, H., He, L., Liu, M., Blaser, M. J., and Li, H. (2018). A two-stage microbial association mapping framework with advanced fdr control. Microbiome, 6(1), 131.

Kaul, A., Mandal, S., Davidov, O., and Peddada, S. D. (2017). Analysis of microbiome data in the presence of excess zeros. Frontiers in microbiology, 8, 2114.

Kleinbaum, D. G., Kupper, L. L., Nizam, A., and Muller, K. G. (2007). Applied Regression Analysis and Other Multivariable Methods. Duxbury Press.

La Rosa, P. S., Brooks, J. P., Deych, E., Boone, E. L., Edwards, D. J., Wang, Q., Sodergren, E., Weinstock, G., and Shannon, W. D. (2012). Hypothesis testing and power calculations for taxonomic-based human microbiome data. PloS one, 7(12), e52078.

Legendre, P. and Anderson, M. J. (1999). Distance-based redundancy analysis: testing multispecies responses in multifactorial ecological experiments. Ecological monographs, 69(1), 1–24.

Love, M. I., Huber, W., and Anders, S. (2014). Moderated estimation of fold change and dispersion for rna-seq data with deseq2. Genome biology, 15(12), 550.

Mandal, S., Van Treuren, W., White, R. A., Eggesbø, M., Knight, R., and Peddada, S. D. (2015). Analysis of composition of microbiomes: a novel method for studying microbial composition. Microbial ecology in health and disease, 26(1), 27663.

McArdle, B. H. and Anderson, M. J. (2001). Fitting multivariate models to community data: a comment on distance-based redundancy analysis. Ecology, 82(1), 290–297.

Morgan, X. C., Kabakchiev, B., Waldron, L., Tyler, A. D., Tickle, T. L., Milgrom, R., Stempak, J. M., Gevers, D., Xavier, R. J., Silverberg, M. S., et al. (2015). Associations between host gene expression, the mucosal microbiome, and clinical outcome in the pelvic pouch of patients with inflammatory bowel disease. Genome biology, 16(1), 67.

Muller, K. E. and Fetterman, B. A. (2012). Regression and ANOVA: An Integrated Approach using SAS Software. SAS Institute.

Paulson, J. N., Stine, O. C., Bravo, H. C., and Pop, M. (2013). Differential abundance analysis for microbial marker-gene surveys. Nature methods, 10(12), 1200–1202.

Robinson, M. D., McCarthy, D. J., and Smyth, G. K. (2010). edger: a bioconductor package for differential expression analysis of digital gene expression data. Bioinformatics, 26(1), 139–140.

Sandve, G. K., Ferkingstad, E., and Nygård, S. (2011). Sequential monte carlo multiple testing. Bioinformatics, 27(23), 3235–3241.

Satten, G. A., Tyx, R. E., Rivera, A. J., and Stanfill, S. (2017). Restoring the duality between principal components of a distance matrix and linear combinations of predictors, with application to studies of the microbiome. PloS one, 12(1), e0168131.

Satten, G. A., Kong, M., and Datta, S. (2018). Multisample adjusted u-statistics that account for confounding covariates. Statistics in Medicine, 37(2), 3357–3372.

Shi, P. and Li, H. (2017). A model for paired-multinomial data and its application to analysis of data on a taxonomic tree. Biometrics, 73(4), 1266–1278.

VanderWeele, T. J. and Shpitser, I. (2011). A new criterion for confounder selection. Biometrics, 67(4), 1406–1413.

Westfall, P. H. and Young, S. S. (1993). Resampling-based multiple testing: Examples and methods for p-value adjustment. John Wiley & Sons.

Wu, C., Chen, J., Kim, J., and Pan, W. (2016). An adaptive association test for microbiome data. Genome medicine, 8(1), 56.

Zhan, X., Tong, X., Zhao, N., Maity, A., Wu, M. C., and Chen, J. (2017). A small-sample multivariate kernel machine test for microbiome association studies. Genetic epidemiology, 41(3), 210–220.

Zhao, N., Chen, J., Carroll, I. M., Ringel-Kulka, T., Epstein, M. P., Zhou, H., Zhou, J. J., Ringel, Y., Li, H., and Wu, M. C. (2015). Testing in microbiome-profiling studies with mirkat, the microbiome regression-based kernel association test. The American Journal of Human Genetics, 96(5), 797–807.

