## Supplemental Materials for "Testing hypotheses about the microbiome using the linear decomposition model (LDM)"

### Supplementary Materials

**Text S1. A variant of the Freedman and Lane permutation scheme**

Here we describe the permutation scheme we use to assess the significance of our test statistics for both the LDM and PERMANOVA-FL. We make use of the fact that, since the columns of $X$ are orthogonal, $X_k X_k^{\mathrm{T}}$ is the orthogonal projection operator (hat matrix) corresponding to variables in submodel $k$. Consider the linear model for the $j$th column of the matrix $Y$ given by

$$Y_{\cdot j} = \sum_{k=1}^{K} X_k \beta_{k;\cdot j} + \epsilon_{\cdot j}, \tag{S1}$$

Suppose we wish to test the $k$th submodel. The Freedman-Lane approach is to form residuals from the reduced model that excludes the term $X_k$. Since the columns of $X$ are orthogonal, we have immediately that residuals for the reduced model are given by

$$Y_{k;\cdot j} = \left( I - \sum_{\substack{k'=1 \\ k' \neq k}}^{K} X_{k'} X_{k'}^{\mathrm{T}} \right) Y_{\cdot j},$$

To generate a new set of values $Y_{\cdot j}^{(\pi)}$ for $Y_{\cdot j}$ in which all linear effects except those corresponding to $X_k$ are preserved, but the residuals are permuted, we write

$$Y_{\cdot j}^{(\pi)} = \left( \sum_{\substack{k'=1 \\ k' \neq k}}^{K} X_{k'} X_{k'}^{\mathrm{T}} \right) Y_{\cdot j} + P_\pi Y_{k;\cdot j},$$

where $P_\pi$ is a permutation matrix. To preserve the correlation structure among OTUs, we use the same permutation matrix $P_\pi$ for each column $j$. In order to construct the $F$ tests we have described, we need to calculate the residuals we would obtain by fitting either the full model (S1) to the permuted data $Y_{\cdot j}^{(\pi)}$, and the reduced model that excludes the term $X_k$. These quantities are most easily obtained by left-multiplying by an appropriate projection operator.

The residual after fitting the full model is given by

$$\left(I - \sum_{k'=1}^{K} X_{k'} X_{k'}^{\mathrm{T}}\right) Y_{\cdot j}^{(\pi)} = \left(I - \sum_{k'=1}^{K} X_{k'} X_{k'}^{\mathrm{T}}\right) P_{\pi} Y_{k;\cdot j}, \tag{S2}$$

so that the residual sum of squares after fitting the full model is

$$Y_{k;\cdot j}^{\mathrm{T}} P_{\pi}^{\mathrm{T}} \left(I - \sum_{k'=1}^{K} X_{k'} X_{k'}^{\mathrm{T}}\right) P_{\pi} Y_{k;\cdot j}. \tag{S3}$$

Because the $X_k$s are orthogonal, the residual and residual sum of squares for the restricted models have the same form as (S2) and (S3) but with sums that are restricted to exclude $k' = k$. Thus, the difference between the residual sum of squares for the full and restricted models is simply the contribution to the sum of squares for $X_k$, given by

$$Y_{k;\cdot j}^{\mathrm{T}} P_{\pi}^{\mathrm{T}} X_k X_k^{\mathrm{T}} P_{\pi} Y_{k;\cdot j}. \tag{S4}$$

Finally, we note that if $P_{\pi}$ is a permutation matrix, then $P_{\pi}^{\mathrm{T}}$ is also a permutation matrix corresponding to the permutation that reverses the effect of $P_{\pi}$, i.e., $P_{\pi} P_{\pi}^{\mathrm{T}} = I$. Thus, we define $X_k^{(\pi)} = P_{\pi}^{\mathrm{T}} X_k$ to be a row-permuted version of $X_k$ and note that the columns of $X^{(\pi)}$ remain orthogonal, so that $X_k^{(\pi)} X_k^{(\pi)^{\mathrm{T}}}$ is the orthogonal projection (hat) matrix corresponding to fitting a model in which the variables have been permuted according to permutation matrix $P_{\pi}^{\mathrm{T}}$. With these observations, we note that (S2) - (S4) can be written entirely in terms of the $X_k^{(\pi)}$s, e.g. (S2) becomes

$$Y_{k;\cdot j}^{\mathrm{T}} \left(I - \sum_{k'=1}^{K} X_{k'}^{(\pi)} X_{k'}^{(\pi)^{\mathrm{T}}}\right) Y_{k;\cdot j},$$

which is the denominator of $F_{kj}$ given in (10) while (S4) becomes

$$Y_{k;\cdot j}^{\mathrm{T}} X_k^{(\pi)} X_k^{(\pi)^{\mathrm{T}}} Y_{k;\cdot j},$$

which is the numerator of $F_{kj}$. Note also that $X_k^{(\pi)^{\mathrm{T}}} X_{k'}^{(\pi)} = \delta_{kk'} I$ since $P_{\pi} P_{\pi}^{\mathrm{T}} = I$.

**Text S2. Simulating read count data with the Poisson log-normal model (PLNM)**

This model assumes that count data for the $j$th OTU of the $i$th observation are generated from independent Poisson distributions with mean $N_i\theta_j$, $j = 1, \ldots, J$, where $N_i$ is a scale factor like the library size, and that the Poisson means $\theta = (\theta_1, \ldots, \theta_J)$ follow a multivariate log-normal distribution with log-mean vector $\mu$ and log-variance-covariance matrix $\Sigma$. In the following, we first describe an approach to estimating these parameters from a real dataset and then outline the steps for simulating read count data using the PLNM.

Let $\mu_j$ denote the $j$th element of $\mu$ and $\sigma_{jk}$ denote the $(j,k)$th element of $\Sigma$. Using results from Aitchison and Ho [1989], the moments of the counts are then given by

$$\mathrm{E}(Y_{ij}) = \exp(\mu_j + \ln(N_i) + 0.5\sigma_{jj}) \equiv N_i\pi_j \;\text{ and }\; \mathrm{Cov}(Y_j, Y_k) = \delta_{jk}N_i\pi_j + N_i^2\pi_j\pi_k\left\{\exp(\sigma_{jk}) - 1\right\},$$

where $\delta_{jk} = 1(j = k)$. The first term in the Covariance is the Poisson variance in the absence of overdispersion (i.e., when $\Sigma = 0$) while the second term is the overdispersion term. It is difficult to obtain the distribution or even moments of $\widehat{\pi}_{ij} = Y_{ij}/\sum_{j'=1}^{J} Y_{ij'}$. Since our goal is merely to choose a reasonable matrix $\Sigma$ to use to generate simulated data, we approximate the moments of $\widehat{\pi}_i = (\widehat{\pi}_{i1}, \ldots, \widehat{\pi}_{iJ})$ by

$$\mathrm{E}(\widehat{\pi}_{ij}) = \pi_j \;\text{ and }\; \mathrm{Cov}(\widehat{\pi}_{ij}, \widehat{\pi}_{ik}) = \delta_{jk}\pi_j - \pi_j\pi_k + \pi_j\pi_k\left\{\exp(\sigma_{jk}) - 1\right\},$$

which correctly accounts for the normalization when $\sigma_{jk} = 0$ by replacing the Poisson variance by the multinomial variance, while retaining the overdispersion term.

Next, suppose we have data on $N$ observations. If $\overline{\pi}$ and $\overline{V}$ are the empirical mean vector and variance-covariance matrix of the $\widehat{\pi}$ values from these observations, the overdispersion (empirical variance minus multinomial variance) is then given by

$$\mathcal{O} = \overline{V} - \mathrm{diag}(\overline{\pi}) + \overline{\pi} \otimes \overline{\pi}.$$

Using the second moment of $\widehat{\pi}$, we form the equation

$$\mathcal{O} = \mathrm{diag}(\overline{\pi})\left\{\exp(\sigma) - 1\right\}\mathrm{diag}(\overline{\pi}),$$

from which we obtain

$$\exp(\sigma) = 1 + \mathrm{diag}(\overline{\pi}^{-1})\mathcal{O}\mathrm{diag}(\overline{\pi}^{-1}),$$

where 1 is a matrix with all elements equal to 1 and $\exp(\sigma)$ is a matrix with the $(i, j)$th element given by $\exp(\sigma_{ij})$. Taking the element-wise logarithm of $\exp(\sigma)$ cannot ensure the positive definiteness of the estimated $\Sigma = \{\sigma_{jk}\}$. We thus make a further approximation and replace the matrix of element-wise exponentials $\exp(\sigma)$ by $\exp(\Sigma)$, the matrix exponential of the variance-covariance matrix. To take the logarithm of a matrix exponential, we first form the eigendecomposition of $\exp(\Sigma)$ by writing

$$\exp(\Sigma) = 1 + \mathrm{diag}(\overline{\pi}^{-1})\mathcal{O}\mathrm{diag}(\overline{\pi}^{-1}) = Q\Lambda Q^T,$$

and then take the logarithm of the eigenvalues to obtain

$$\Sigma = Q_r(\ln \Lambda_r)Q_r^T,$$

where the subscript $r$ refers to the restriction to positive values of $\Lambda$ and columns of $Q$ corresponding to positive values of $\Lambda$. Once we have obtained $\Sigma$, we can find $\mu$ by solving the equations

$$\exp(\mu_j + 0.5\sigma_{jj}^2) = \overline{\pi}_j.$$

We estimated $\mu$ and $\Sigma$ from the URT (i.e., upper-respiratory-tract) data and generated the read count data as follows. We generated a set of Poisson means $\theta$ from the log-normal distribution with log-mean $\mu$ and log-variance-covariance matrix $\Sigma$; we scaled the $\theta$ values by the appropriate library size, and generated an independent Poisson random count for each OTU. We first used this method to simulate data uniformly for cases and controls. The data for cases were treated as the "baseline" (or "initial") data. We then created "disease" data for cases by shuffling the "baseline" data among the OTUs selected for $U$ (the case-control status) in S1 or S2. The final data for cases were chosen between the "disease" and "baseline" data with probabilities $\beta$ and $1 - \beta$, where $\beta$ represents the strength of association. Note that no confounders were generated in the simulation using the PLNM.

**Text S3. Simulating read count data with the Negative-Binomial (NB) model**

Simulations using the NB model were based on the Dirichlet-Multinomial model simulations for Scenarios S1 and S2. As in the main text, we used the one-way, case-control design with a confounder and 100 independent samples. The only difference was that read count data were generated from the NB model. For the read count data for the $i$th sample at the $j$th OTU, we chose the mean parameter of the NB model to be the $j$th element of the sample-specific frequency vector $\widetilde{\pi}(U_i, C_i)$ (defined in the main texts) multiplied by the simulated library size for the $i$th sample. We chose the overdispersion parameter of the NB model to be the estimated overdispersion from fitting the NB model to data from the $j$th OTU of the real throat data. We verified that the mean and variance of the NB-simulated count data at each OTU matched well with the real data; also, the overall proportion of zero counts matched well with the real data.

**Text S4. Simulating clustered data, data with a two-way design, and data with a quantitative trait**

To generate clustered data, we assume that we had samples from 50 distinct individuals, each of whom contributed 2 samples. We modified scenarios S1 and S2 by assuming that half of the individuals were cases ($U = 1$) and the remaining individuals were controls ($U = 0$). We generated the confounder $C$ at the individual level in the same way as for unclustered data; to induce within-cluster correlations, we generated individual-specific OTU frequencies from the Dirichlet distribution with mean frequencies $\widetilde{\pi}(U, C)$ (defined in the main texts) and overdispersion parameter 0.02, and then generated counts for each samples from the same individual using the Multinomial distribution with mean being the individual-specific OTU frequencies, using library sizes that were generated independently for each sample. Note that if each individual had a single sample, the combination of Dirichlet and Multinomial sampling would reproduce the DM mixture model used for unclustered data.

To simulate data with a two-way design (without confounding), we considered two factors, $U_1$ and $U_2$, that each have two levels and that are orthogonal. The samples were randomly split into two groups with equal size, one group being assigned $U_1 = 0$ and the other $U_1 = 1$.

Samples in each group were then further split randomly into two subgroups with equal size, with one group assigned $U_2 = 0$ and the other $U_2 = 1$. Then we induced association of $U_1$ and $U_2$ with the OTUs in the same way as $U$ and $C$ by assigning $U_1$ and $U_2$ the same sets of OTUs as assigned to $U$ and $C$ in scenarios S1 and S2 and using the sample sample-specific frequency vector $\widetilde{\pi}(U_1, U_2) = (1 - \beta_1 U_1 - \beta_2 U_2)\pi_1 + \beta_1 U_1 \pi_2 + \beta_2 U_2 \pi_3$. Note that $\beta_1$ and $\beta_2$ are the effect sizes of $U_1$ and $U_2$, respectively.

To generate data with a continuous trait, we used a model considered by Zhao *et al.* [2015]. We first generated OTU counts for each sample using the DM model with frequency vector $\pi_1$, overdispersion parameter 0.02, and library size sampled from $N(10000, 10000/3)$. Let $S = \sum_{j \in \mathcal{A}} Y_{ij}/\overline{Y}_j$, where $\mathcal{A}$ is the set of the ten most abundant OTUs, $Y_{ij}$ is the frequency of the $j$th OTU in the $i$th sample, and $\overline{Y}_j$ is the average frequency for the $j$th OTU across samples. We generated a confounder $C = \text{scale}(S) + \widetilde{\epsilon}$, where $\text{scale}(v)$ centers and normalizes vector $v$ to have unit variance and $\widetilde{\epsilon} \sim N(0, 1)$. Finally, we simulated the continuous trait as $U = \beta_C \text{scale}(C) + \beta \text{scale}(S) + \epsilon$, where $\beta_C = 0.3$ and $\epsilon \sim N(0, 1)$. Note that when $\beta = 0$ there is no association between $U$ and the OTU frequencies.

**Text S5. The LDM and Redundancy Analysis**

The LDM bears some resemblance to Redundancy Analysis (RA), but also differs in notable respects. RA seeks to describe how much of a matrix $Y$ can be explained by a single set of variables $X_1$, also concluding that the variability explained is $||X_1 X_1^{\mathrm{T}} Y||_F^2$. RA also calculates a matrix like $\beta_1$; however, RA requires that $\beta_1$ have orthogonal columns, which is unnecessary for calculating either $\mathrm{Tr}(D_1^2)$ or $|\beta_{1;\cdot j}|^2$. Further, RA only allows analysis of one set of variables at a time, so only a single matrix $\beta_1$ is produced; this is presumably because the non-orthogonality of multiple $\beta_k$s implies that it is impossible to find $\beta_1$ and $\beta_2$ that satisfy $\beta_1 \beta_2^{\mathrm{T}} = 0$ for arbitrary submodels $X_1$ and $X_2$. Thus in RA, the effect of each submodel $X_k$ must be tested sequentially using a separate linear model like

$$\widetilde{Y}_k = X_k \beta_k + \epsilon,$$

where

$$\widetilde{Y}_k = \left( I - \sum_{k' < k} X_{k'} X_{k'}^{\mathrm{T}} \right) Y.$$

As a result, the $F$ tests available in the LDM are expected to be more powerful than the type I or "order of variables added" tests available in RA when there is more than one submodel [Muller and Fetterman, 2012]. This is because the residual sums of squares in the denominator of the type III tests used in the LDM include *all* submodels tested, rather than only submodels with $k' < k$ used in sequential RA. Use of the restricted model in RA can thus result in an incorrect estimate of the residual sum of squares, which may affect power even in a permutation setting as the test is then not (asymptotically) pivotal. A second advantage of the LDM is that it is that we can assign significance to all submodels with a single permutation experiment, while RA requires a separate set of permutations for each submodel $X_k$ tested.

#### References


Aitchison, J. and Ho, C. (1989). The multivariate poisson-log normal distribution. *Biometrika*, **76**(4), 643–653.

Muller, K. E. and Fetterman, B. A. (2012). *Regression and ANOVA: An Integrated Approach using SAS Software*. SAS Institute.

Zhao, N., Chen, J., Carroll, I. M., Ringel-Kulka, T., Epstein, M. P., Zhou, H., Zhou, J. J., Ringel, Y., Li, H., and Wu, M. C. (2015). Testing in microbiome-profiling studies with mirkat, the microbiome regression-based kernel association test. *The American Journal of Human Genetics*, **96**(5), 797–807.

**Table S1** Type I error for testing the global hypothesis when read count data were generated from the Poisson log-normal model (PLNM)

| Scenario | PERMANOVA-FL | VE-freq | VE-arcsin | VE-omni |
|---|---|---|---|---|
| S1 and S2 | 0.051 | 0.046 | 0.053 | 0.051 |

All methods adjusted for the confounder. PERMANOVA-FL was based on the Bray-Curtis distance. Scenarios S1 and S2 are equivalent when the effect size $\beta$ is zero.

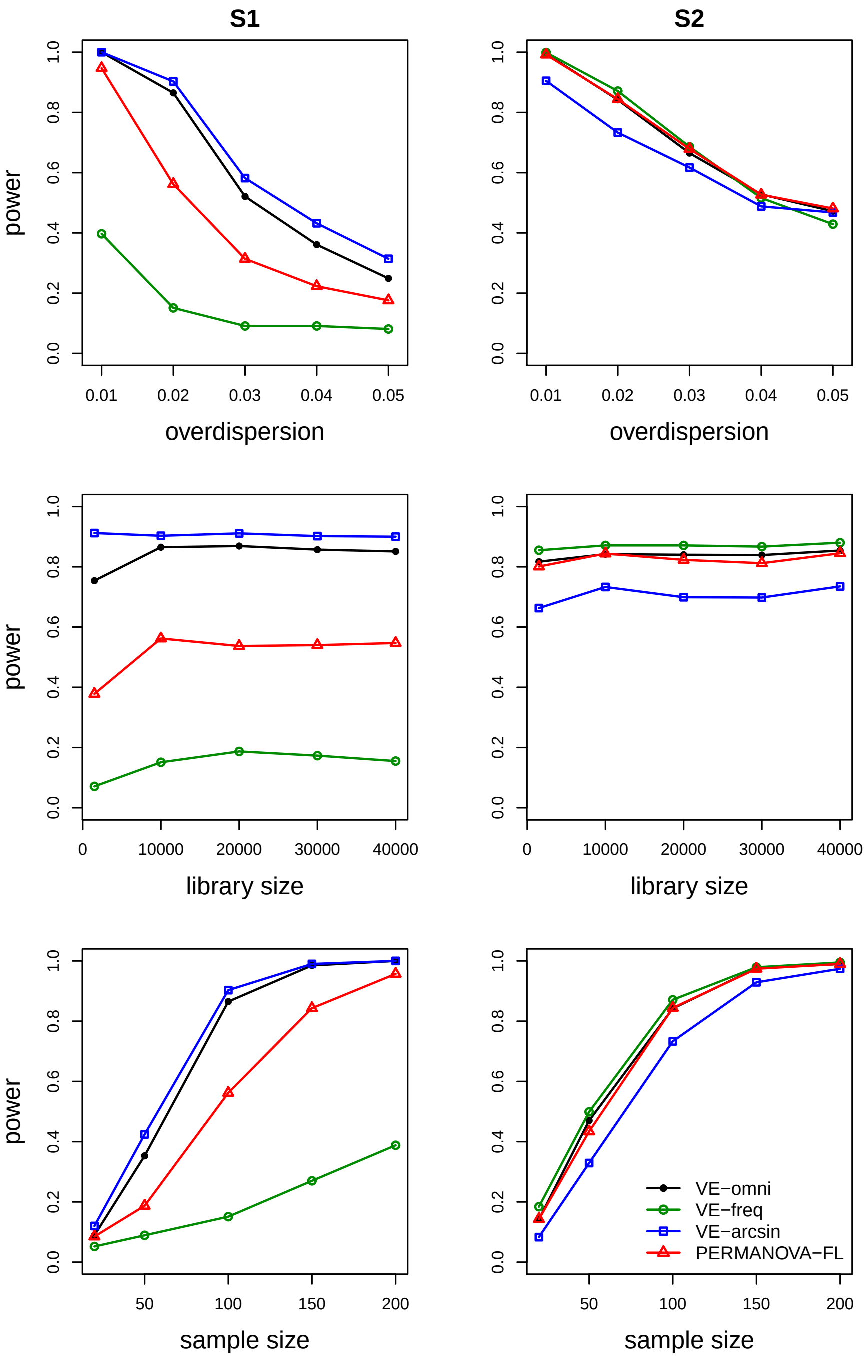


**Fig. S1** Power for testing the global hypothesis with different overdispersion values (top panel), library sizes (middle panel), and sample sizes (bottom panel). At each mean library size $\mu$, the individual library size was sampled from $N(\mu, \mu/3)$ and left-truncated at 500. When not otherwise stated, the over-dispersion parameter is 0.02, mean library size is 10000, and sample size is 100. The effect size $\beta$ is 0.14 for S1 and 0.35 for S2. PERMANOVA-FL was based on the Bray-Curtis distance.

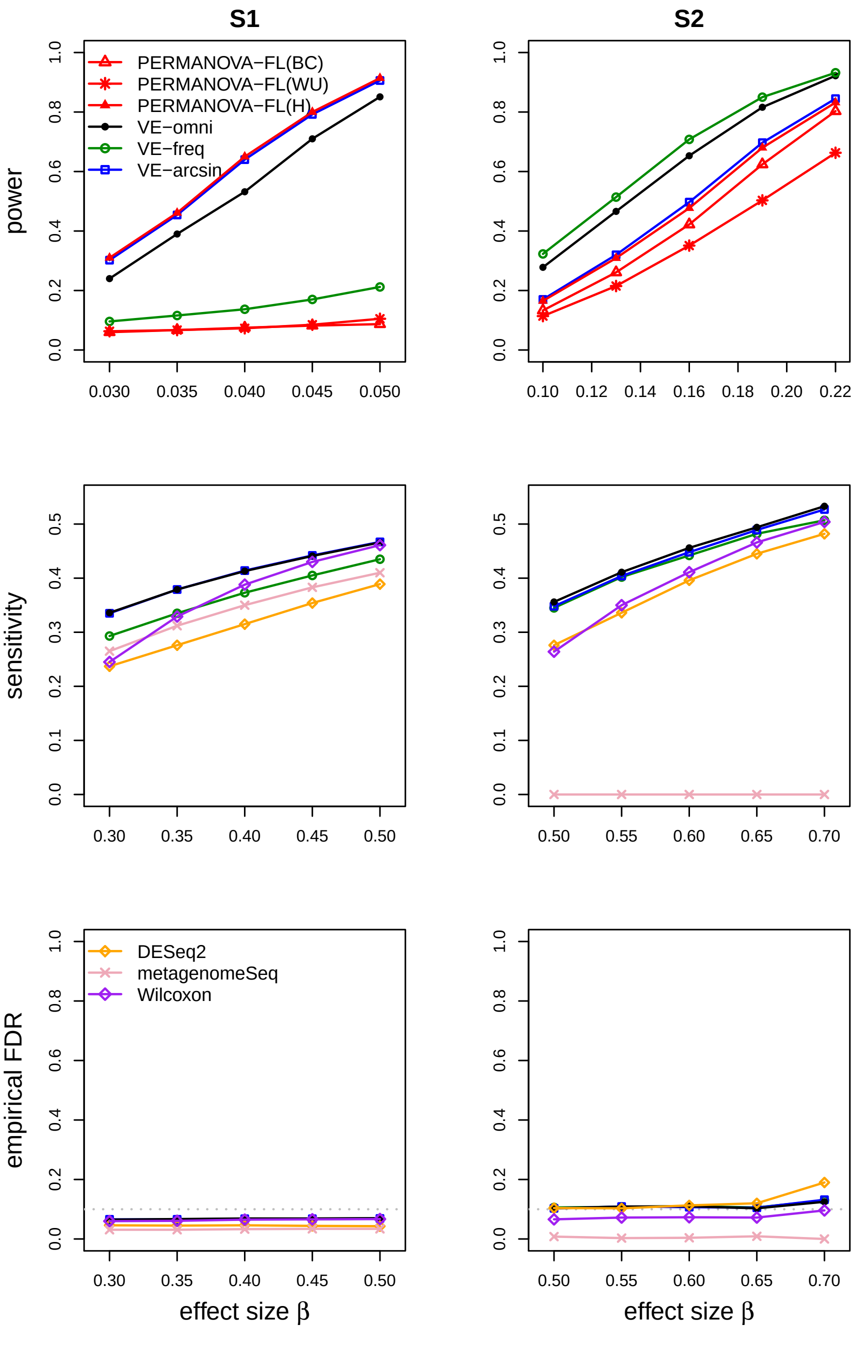


**Fig. S2** Simulation results when the read count data were generated by the PLNM. The gray dotted lines represent the nominal FDR 0.1. BC: Bray-Curtis; WU: weighted UniFrac; H: Hellinger. MetagenomeSeq and the Wilcoxon test were applied because of the absence of confounders in the simulation using the PLNM.

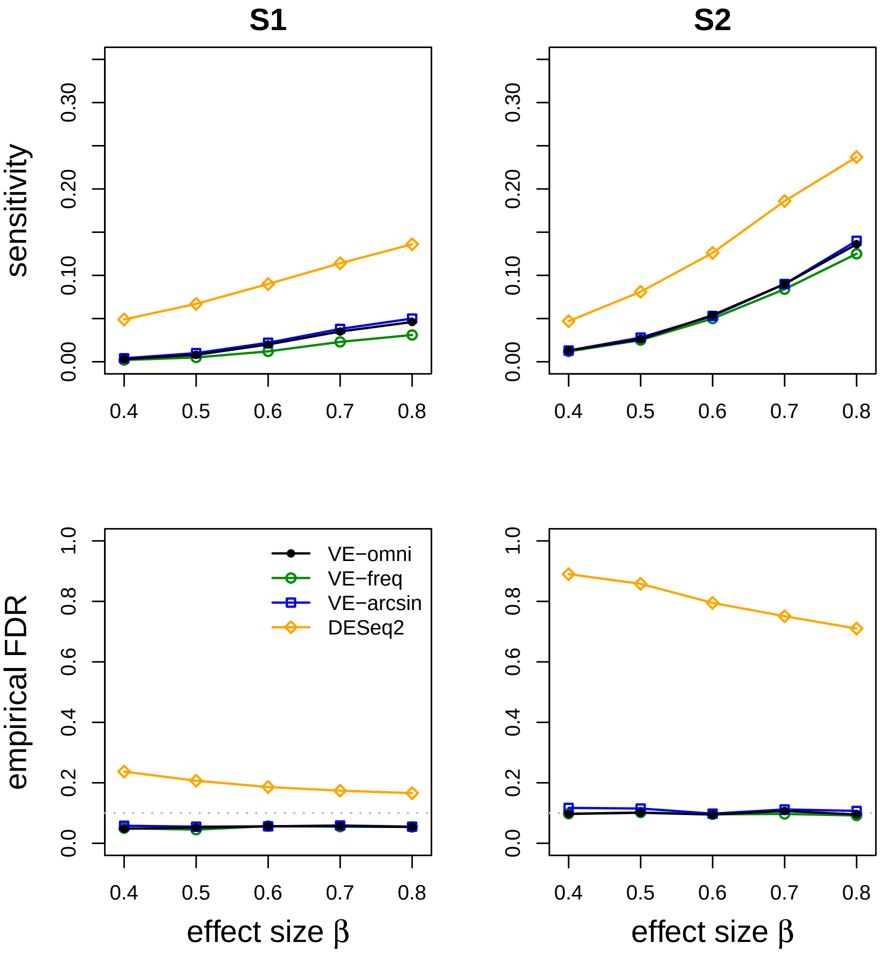


**Fig. S3** Sensitivity and empirical FDR for testing differentially abundant OTUs when the read count data were generated by the NB model. The gray dotted lines represent the nominal FDR 0.1.

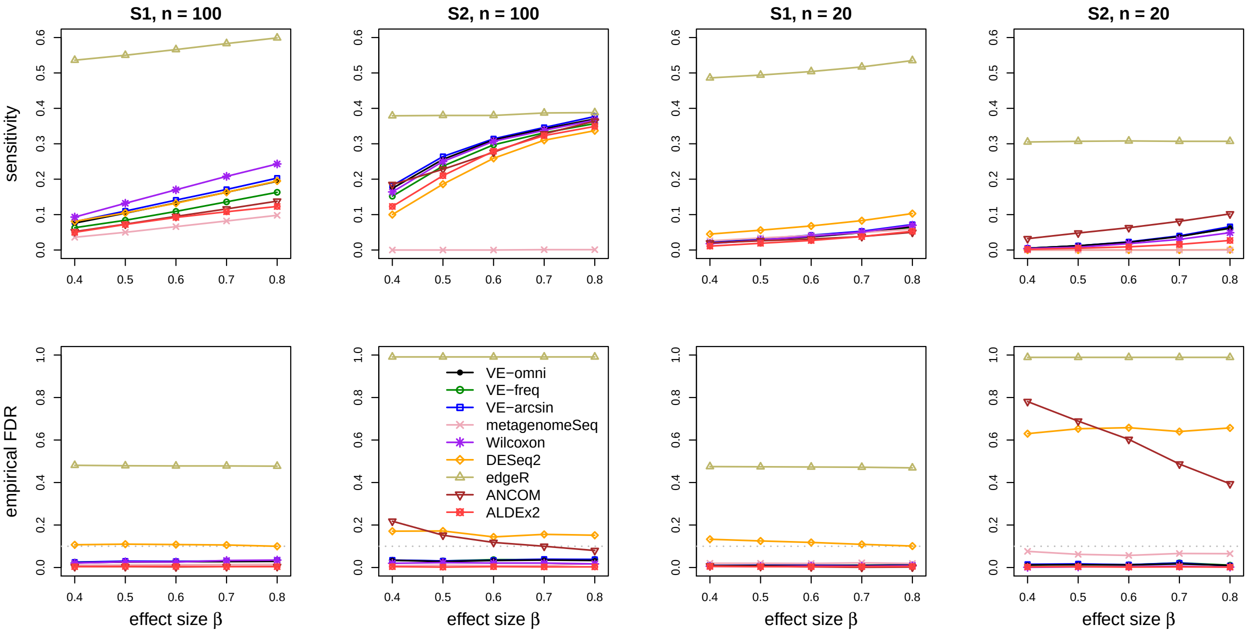


**Fig. S4** Sensitivity and empirical FDR for testing differentially abundant OTUs in absence of confounders. The gray dotted lines represent the nominal FDR 0.1.

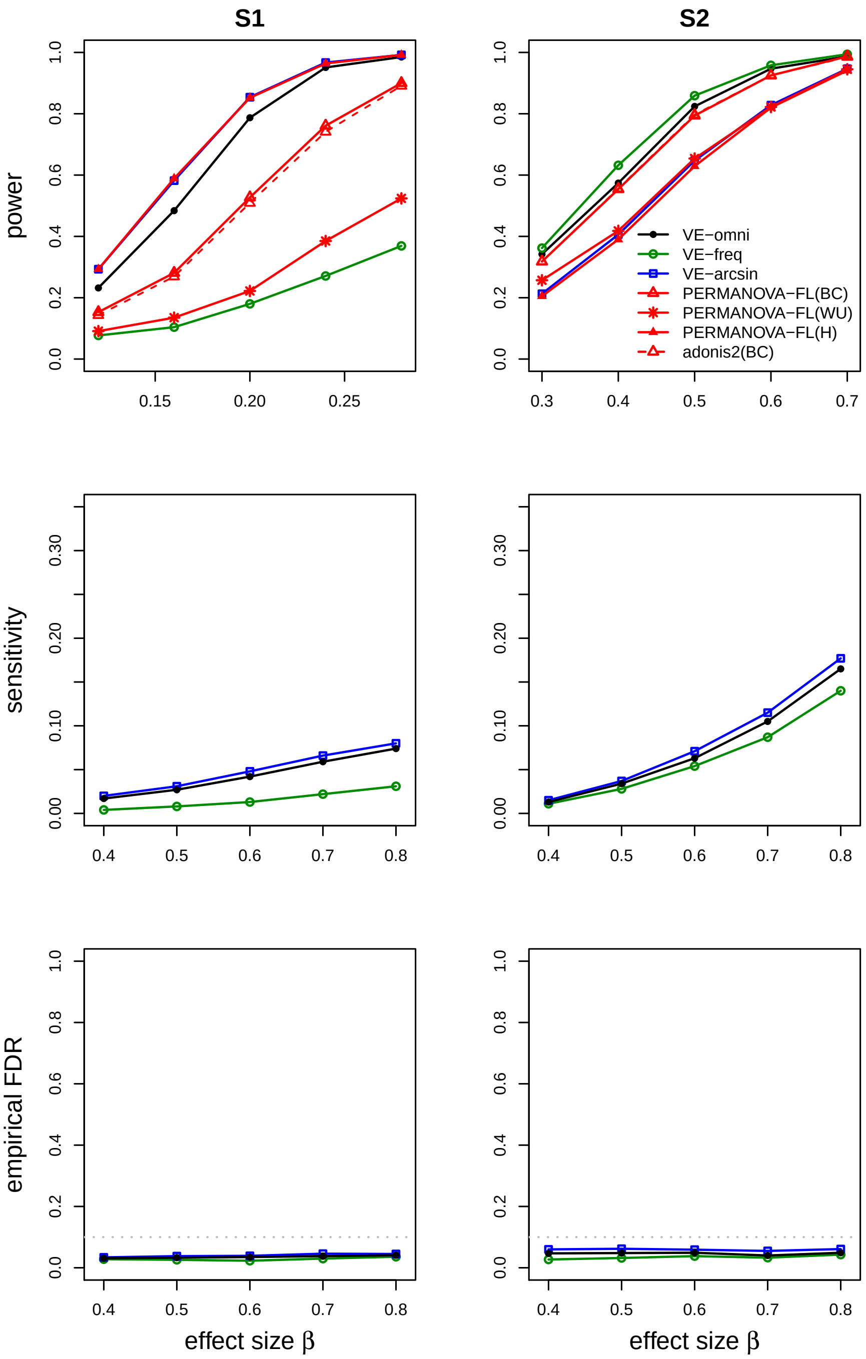


**Fig. S5** Simulation results for clustered data. The gray dotted lines represent the nominal FDR=0.1. BC: Bray-Curtis; WU: weighted UniFrac; H: Hellinger. The DESeq2 program is not applicable for this type of clustered data.

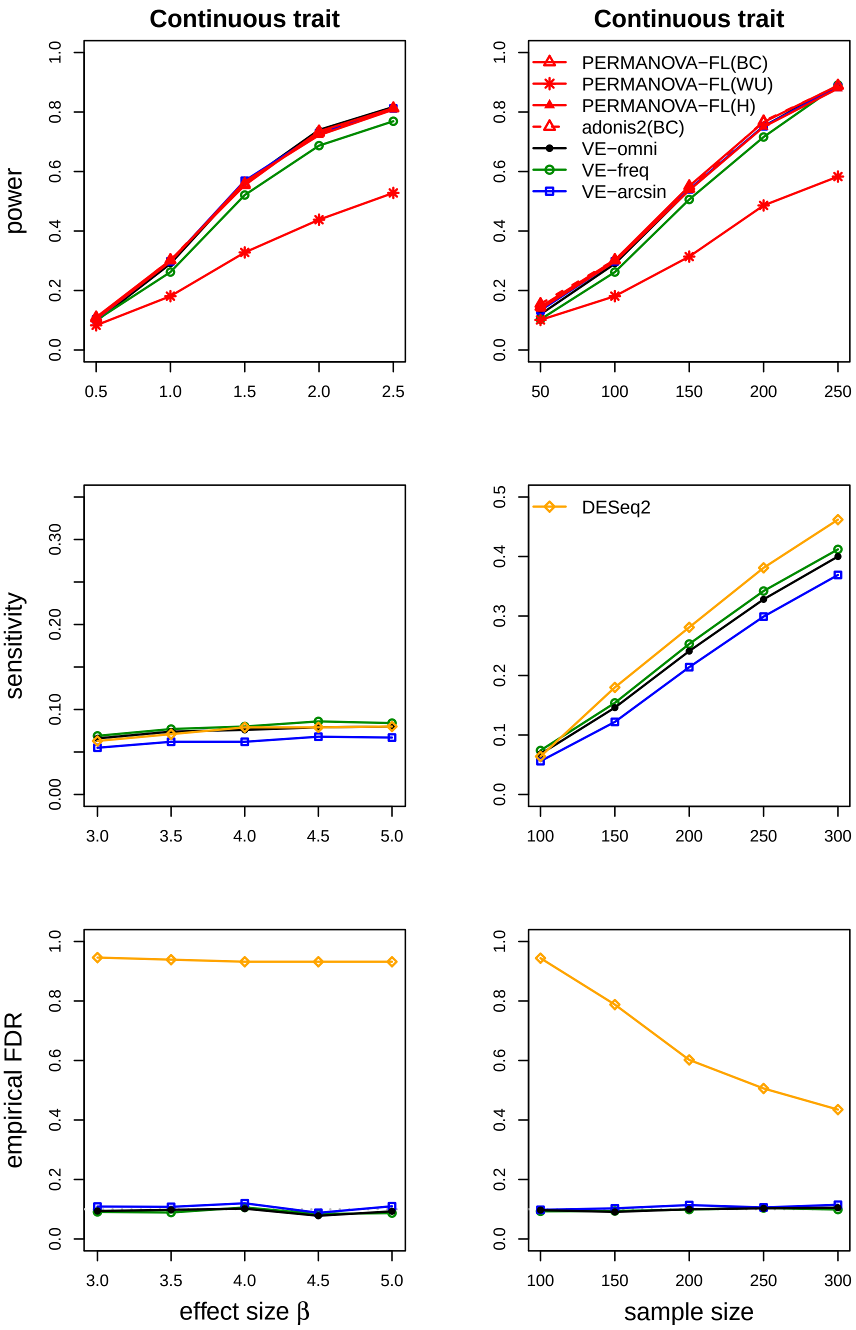


**Fig. S6** Simulation results for a continuous trait. The first and second columns correspond to results as the effect size and the sample size, respectively, increase. The gray dotted lines represent the nominal FDR=0.1. When varying the sample size, we set $\beta = 1$ for evaluating power and $\beta = 3$ for sensitivity and empirical FDR.

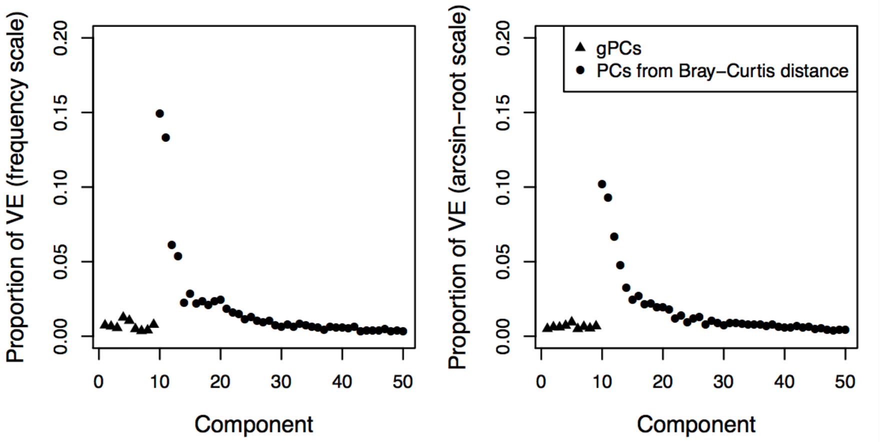


**Fig. S7** Exploratory analysis of the PPI microbiome data based on the Bray-Curtis distance. The proportions of variance explained by the 9 gPCs and the PCs of the (residual) distance measure are obtained after removing the effects of confounders antibiotic use, inflammation score, and disease type. The PCs are ordered by their Bray-Curtis eigenvalues. The left plot is based on frequency data and the right plot is based on arcsin-root transformed data. The components are ordered by the magnitude of their corresponding eigenvalue in a spectral decomposition of $\Delta_{10}$ (the distance matrix after removing the effect of confounders and the 9 gPCs). Only the first 50 (of 195 total) components are shown.
